# The parabrachial to central amygdala circuit is a key mediator of injury-induced pain sensitization

**DOI:** 10.1101/2023.02.08.527340

**Authors:** Jeitzel M. Torres-Rodriguez, Torri D. Wilson, Sudhuman Singh, Sarah Chaudhry, Anisha P. Adke, Jordan J. Becker, Jenny L. Lin, Santiago Martinez Gonzalez, Omar Soler-Cedeño, Yarimar Carrasquillo

**Author notes:** authors contributed equally (Jeitzel M. Torres-Rodriguez, Torri D. Wilson). Correspondence, Yarimar Carrasquillo, PhD National Institutes of Health 35 Convent Drive, Building 35A / Room 1E-410 Bethesda, MD 20892. **Author Contributions:** Conceptualization, J.T.R., T.D.W., A.A., S.C., O.S.C., Y.C.; Investigation, T.D.W., J.T.R., S.S., S.C., A.A., J.J.B., J.L.L., S.M.G., O.S.C. Data Analysis, J.T.R., T.D.W., S.C., A.A., J.J.B., J.L.L., O.S.C., Y.C.; Writing, J.T.R., T.D.W., S.C., J.L.L., Y.C.; Funding Acquisition, Y.C., S.S., O.S.C. **Competing Interest Statement:** The authors declare no conflict of interests.

## Abstract

The spino-ponto-amygdaloid pathway is a major ascending circuit relaying nociceptive information from the spinal cord to the brain. Potentiation of excitatory synaptic transmission in the parabrachial nucleus (PbN) to central amygdala (CeA) pathway has been reported in rodent models of persistent pain. At the behavioral level, the PbN→CeA pathway has been proposed to serve as a general alarm system to potential threats that modulates pain-related escape behaviors, threat memory, aversion, and affective-motivational (but not somatosensory) responses to painful stimuli. Increased sensitivity to previously innocuous somatosensory stimulation is a hallmark of chronic pain. Whether the PbN→CeA circuit contributes to heightened peripheral sensitivity following an injury, however, remains unknown. Here, we demonstrate that activation of CeA-projecting PbN neurons contributes to injury-induced behavioral hypersensitivity but not baseline nociception in male and female mice. Using optogenetic assisted circuit mapping, we confirmed a functional excitatory projection from PbN→CeA that is independent of the genetic or firing identity of CeA cells. We then showed that peripheral noxious stimulation increases the expression of the neuronal activity marker c-Fos in CeA-projecting PbN neurons and chemogenetic inactivation of these cells reduces behavioral hypersensitivity in models of neuropathic and inflammatory pain without affecting baseline nociception. Lastly, we show that chemogenetic activation of CeA-projecting PbN neurons is sufficient to induce bilateral hypersensitivity without injury. Together, our results demonstrate that the PbN→CeA pathway is a key modulator of pain-related behaviors that can amplify responses to somatosensory stimulation in pathological states without affecting nociception under normal physiological conditions.

**Significance Statement:** Early studies identified the spino-ponto-amygdaloid pathway as a major ascending circuit conveying nociceptive inputs from the spinal cord to the brain. The functional significance of this circuit to injury-induced hypersensitivity, however, remains unknown. Here, we addressed this gap in knowledge using viral-mediated anatomical tracers, *ex-vivo* electrophysiology and chemogenetic intersectional approaches in rodent models of persistent pain. We found that activation of this pathway contributes to injury-induced hypersensitivity, directly demonstrating a critical function of the PbN→CeA circuit in pain modulation.

## Introduction

Chronic pain is a multidimensional experience that encompasses reflexive-defensive somatosensory and affective-motivational components. In the United States alone, more than 1 in 5 Americans suffer from this condition (1), with similar statistics observed across 52 countries (2). Despite this, diagnostic tools are limited and currently available treatments remain ineffective and addictive (3). These statistics underscore the importance of identifying mechanisms involved in pain processing to potentially improve diagnosis and develop prospective treatments.

The spino-ponto-amygdaloid pathway has been characterized as a major ascending pathway involved in the relay of nociceptive information from the spinal cord to the brain (4–7). In this pathway, peripheral nociceptors receive and relay nociceptive inputs to second order neurons in lamina I of the spinal cord, which then send projections to the pontine parabrachial nucleus (PbN) (8–11). Multiple studies have demonstrated that PbN neurons respond to variety of noxious stimuli (8, 12–14) and function as a hub for the relay of nociceptive information to multiple brain regions, including the periaqueductal gray, hypothalamic and thalamic nuclei, the extended amygdala, and the central amygdala (CeA) (12, 15–17).

Among these brain structures, the CeA is anatomically well-positioned to integrate somatosensory and affective signals within the brain. It receives somatosensory signals via the spino-ponto-amygdaloid pain pathway and polymodal information, including those related to affective and cognitive states, via inputs from the basolateral and lateral amygdala nuclei, which in turn receive inputs from cortical and thalamic regions (18). *In-vivo* and *ex-vivo* studies have further shown that CeA neurons respond to peripheral noxious stimuli and are sensitized following injury in various rodent models of pain (19–21). At the behavioral level, manipulations of CeA neurons have been shown to modulate reflexive-defensive and affective-motivational components of pain in response to injury as well as to mediate different types of analgesia (22–26).

The proposed function of the PbN→CeA pathway in pain processing is further supported by *ex-vivo* electrophysiological studies that have shown potentiation of glutamatergic synaptic transmission in response to injury (20, 21, 27–30). While previous behavioral studies showed that CeA-projecting PbN neurons modulate escape behaviors, affective-motivational responses to painful stimuli, aversion, and threat memory (12, 31, 32), the functional contribution of this circuit to injury-induced hypersensitivity remains unknown. In the present study, we evaluated the function of the PbN→CeA pathway in baseline nociception and injury-induced hypersensitivity using mouse models of inflammatory and neuropathic pain coupled with viral-mediated anatomical tracers, optogenetic assisted circuit mapping, and intersectional chemogenetic strategies. Our results demonstrate that the PbN→CeA pathway is both necessary and sufficient to modulate reflexive-defensive somatosensory responses in pathological states, without altering baseline nociception.

## Results

### Excitatory projection from PbN→CeA is independent of the genetic or firing identity of CeA cells

To validate the functional circuitry between the PbN and CeA, we used optogenetically assisted circuit mapping in acute brain slices from *Sst*-cre::Ai9 or *Prkcd*-cre::Ai9 mice injected with AAV-hChR2-EYFP into the PbN (**Figure 1A-C**). Consistent with previous reports (4, 27) fluorescent terminals were readily observed within the laterocapsular subdivision of the CeA (CeLC) when the Channelrhodopsin-2 (ChR2)-expressing virus was injected into the PbN (**Figure 1B**). Consistent with previous reports, firing phenotypes in PKCδ+ and Som+ CeA neurons were heterogeneous (33–35) (**Figure 1D**). Blue light stimulation of PbN terminals in the CeA evoked excitatory postsynaptic currents (oEPSCs) in the majority of PKCδ+ (80%) and Som+ neurons (90%) recorded (**Figure 1E-F**). These results were true regardless of firing phenotype (**Figure 1E**). Together, these findings confirm that CeLC neurons receive excitatory efferent projections originating from the PbN independently of the genetic or firing identity of CeA neurons.

**Figure 1:**
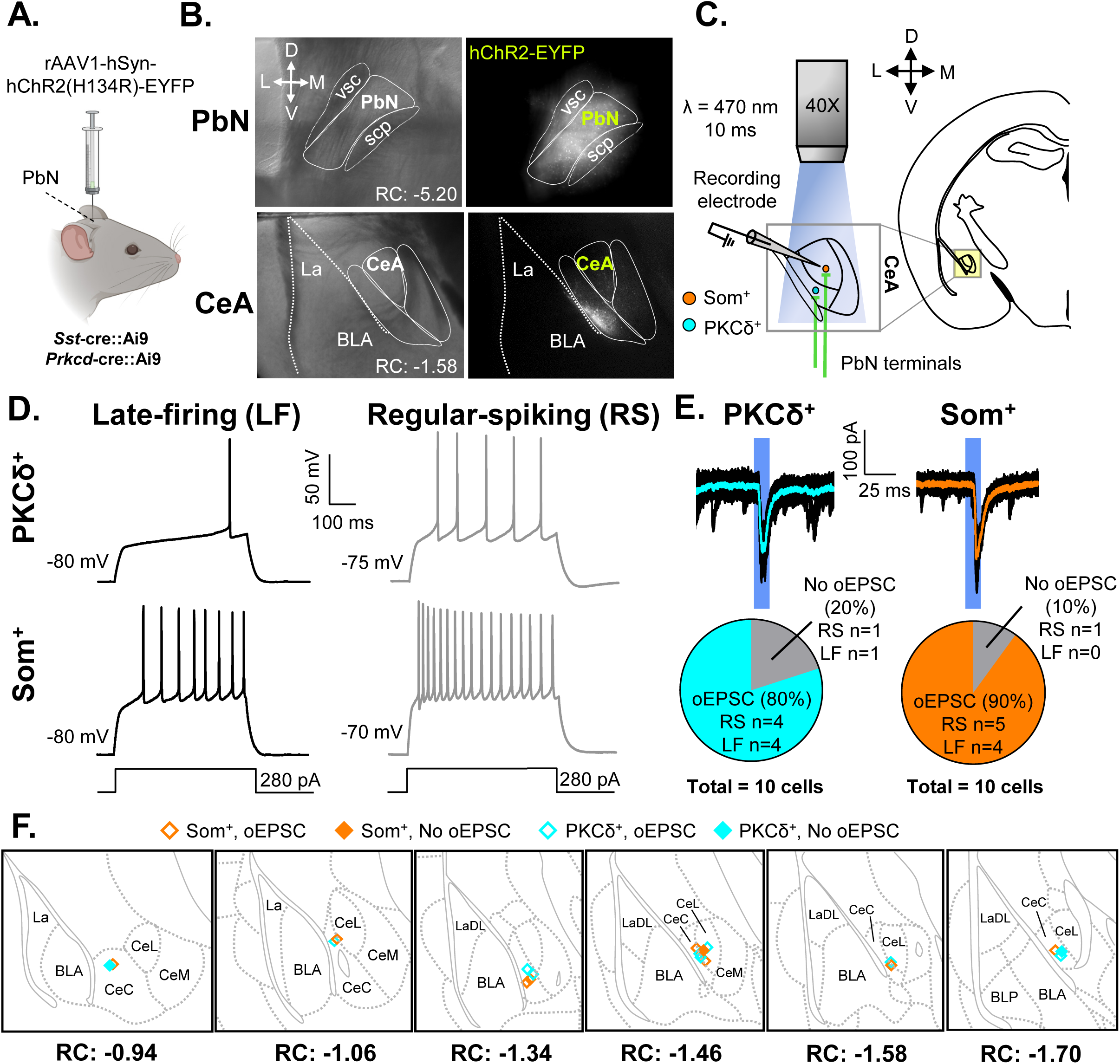
Validation of functional projection from PbN to CeA. **(A)** Male Som-Cre::Ai9 and PKCδ::Ai9 mice were stereotaxically injected with rAAV1-hSyn-hChR2(H134R)-EYFP into the PbN. Whole-cell patch-clamp recordings were performed on mice at least 4 weeks following the injection. **(B)** Representative images of the PbN (top) and CeA (bottom). Left panel depicts differential interference contrast images. Right panel depicts fluorescent images of transduced cells within the PbN and fluorescent PbN terminals within the CeA. **(C)** Schematic depicting optogenetic assisted circuit mapping experiments. Whole-cell patch-clamp recordings of optically evoked excitatory postsynaptic current (oEPSC) in response to blue light (470 nm) stimulation of PbN terminals in CeA-Som+ or CeA-PKCδ+ neurons. **(D)** Representative voltage traces of late-firing (LF; left panel) and regular-spiking (RS; right panel) CeA-PKCδ+ (top) and CeA-Som+ (bottom) neurons in response to a 280-pA depolarizing current injection. **(E)** Representative current traces of oEPSCs in CeA-PKCδ+ and CeA-Som+ neurons in response to a 10 ms pulse of blue light stimulation. The proportion of CeA-PKCδ+ LF and RS neurons displaying oEPSCs was comparable to the proportion of CeA-Som+ LF and RS neurons displaying oEPSCs. **(F)** Recording site illustrations of CeA-Som+ and CeA-PKCδ+ neurons classified by either the presence or absence of oEPSCs. Each symbol highlights the location of an individual neuron. Abbreviations: ventral spinocerebellar tract (vsc), superior cerebellar peduncle (scp), parabrachial nucleus (PbN), central amygdala (CeA), central amygdala medial (CeM), central amygdala lateral (CeL), central amygdala capsular (CeC), basolateral amygdaloid (BLA), lateral amygdaloid (LA).

### Peripheral noxious stimulation induces c-Fos expression in CeA-projecting PbN neurons

Previous studies have shown that CeA-projecting PbN neurons do not receive direct nociceptive inputs from the spinal cord (7, 36, 37). Separate studies have demonstrated that the PbN is activated in response to peripheral noxious stimulation (12, 13, 38, 39) and that injury potentiates the PbN→CeA pathway in several mouse models of persistent pain. Whether CeA-projecting PbN neurons are activated by peripheral noxious stimulation remains unknown. To evaluate this, we measured c-Fos, a surrogate for neuronal activation, in response to pinch stimulation of the hind paw. An intersectional genetic approach was used for the identification of CeA-projecting PbN neurons. C57BL/6J mice were injected with a cre-expressing retrograde Adeno-associated virus (AAV) (pENN.AAV.hSyn.HI.eGFP-Cre.WPRE.SV40) into the CeA and a cre-dependent AAV encoding the red fluorescent protein mCherry (AAV8-hSyn-DIO-mcherry) into the PbN (**Figure 2A**). A pAAV.CMV encoding LacZ was co-injected with the cre-expressing retrograde AAV to identify the injection site in the CeA. As illustrated in **Figure 2B-D**, LacZ expression was restricted to the CeA, and robust transduction efficacy was observed at all CeA rostro-caudal levels. Evaluation of mCherry expression in the PbN showed robust transduction throughout the rostro-caudal PbN that was mostly restricted to the external lateral PbN (**Figure 2E-G**). These results confirm that we can selectively visualize CeA-projecting PbN neurons using this intersectional genetic strategy.

**Figure 2:**
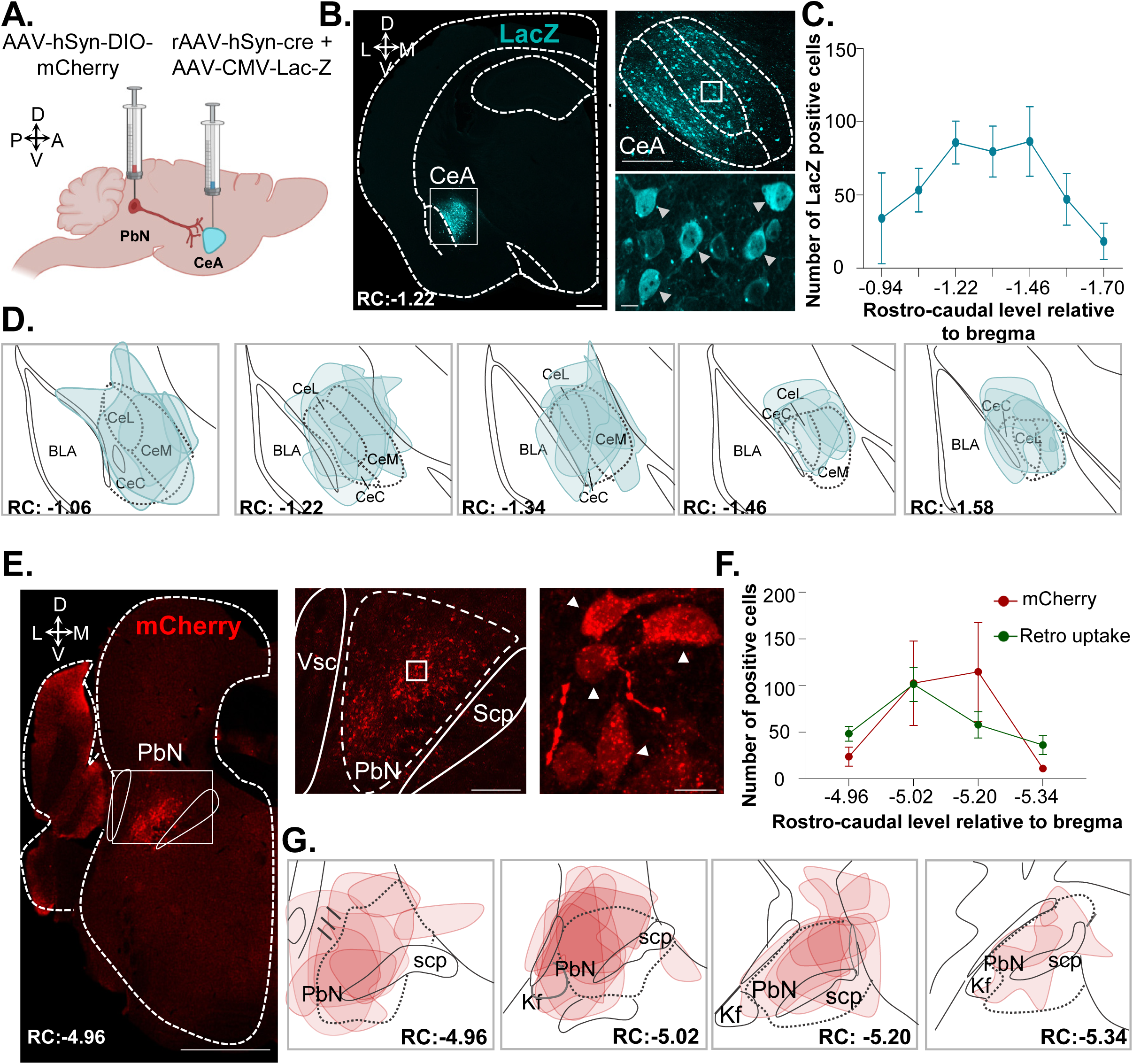
Validation of intersectional genetic approach. **(A)** Schematic of intersectional approach. Male C57BL/6J mice were stereotaxically co-injected with AAV8-hSyn-DIO-mcherry into the right PbN and with a mix (1:1) of AAV.hSyn.HI.eGFP-Cre and pAAV.CMV.LacZ.bGH into the right CeA. **(B)** Representative images of the pAAV.CMV.LacZ.bGH injection into the CeA in a coronal brain slice. Immunofluorescence for LacZ is shown in cyan. Left panel shows low magnification image. Scale bar represents 500 µm. Right panel shows high magnification images of area delineated by the white box. Scale bar for the top panel represents 200 µm and 10 µm for the bottom. Arrows represent transduced cells. Rostro-caudal level relative to bregma (40) is -1.22. **(C)** Mean ± SEM number of LacZ transduced cells by rostro-caudal distribution (n=4 mice and four to six slices per mouse). **(D)** Drawings showing the rostro-caudal distribution of the pAAV.CMV.LacZ.bGH injection into the CeA. **(E)** Representative images of the AAV8-hSyn-DIO-mcherry injection into the PbN in a coronal brain slice. Immunofluorescence for mCherry is shown in red. Left panel shows low magnification image. Scale bar represents 500 µm. Middle and left panels show high magnification images of area delineated by white box. Arrows represent transduced cells. Scale bar for the middle panel represents 200 µm and 10 µm for the left. RC is -4.96. **(F)** Mean ± SEM number of cells with retrograde uptake following AAV.hSyn.HI.eGFP-Cre injection and mCherry-transduced cells by RC (n=10 mice for mCherry and n=25 mice for retrograde uptake; 2-4 slices per mouse). **(G):** Drawings showing the rostro-caudal distribution of the AAV8-hSyn-DIO injection into the PbN. Abbreviations: superior cerebellar peduncle (scp), parabrachial nucleus (PbN), central amygdala (CeA), central amygdala medial (CeM), central amygdala lateral (CeL), central amygdala capsular (CeC), basolateral amygdaloid (BLA).

Consistent with previous studies (12, 13), evaluation of c-Fos expression in the PbN of male and female mice showed significant (p < 0.001) increases in c-Fos+ neurons in response to pinch stimulation of the hind paw, when compared to control mice that did not receive pinch (**Figure 3A-3C**). Quantification of mCherry+ CeA-projecting PbN neurons co-expressing c-Fos further showed that approximately 10% of pinch-induced c-Fos is localized to CeA-projecting PbN neurons (**Figure 3D**) and approximately 20% of CeA-projecting PbN neurons express c-Fos after pinching (**Figure 3E**). In contrast, significantly (p < 0.05) lower co-expression of c-Fos is seen in CeA-projecting PbN neurons in control no-pinch conditions. No overt sex differences in number of positive cells were observed in any of the groups evaluated (**Figure 3D-E**). These combined results demonstrate that pinch stimulation of the hind paw elicits c-Fos expression in CeA-projecting PbN neurons.

**Fig 3:**
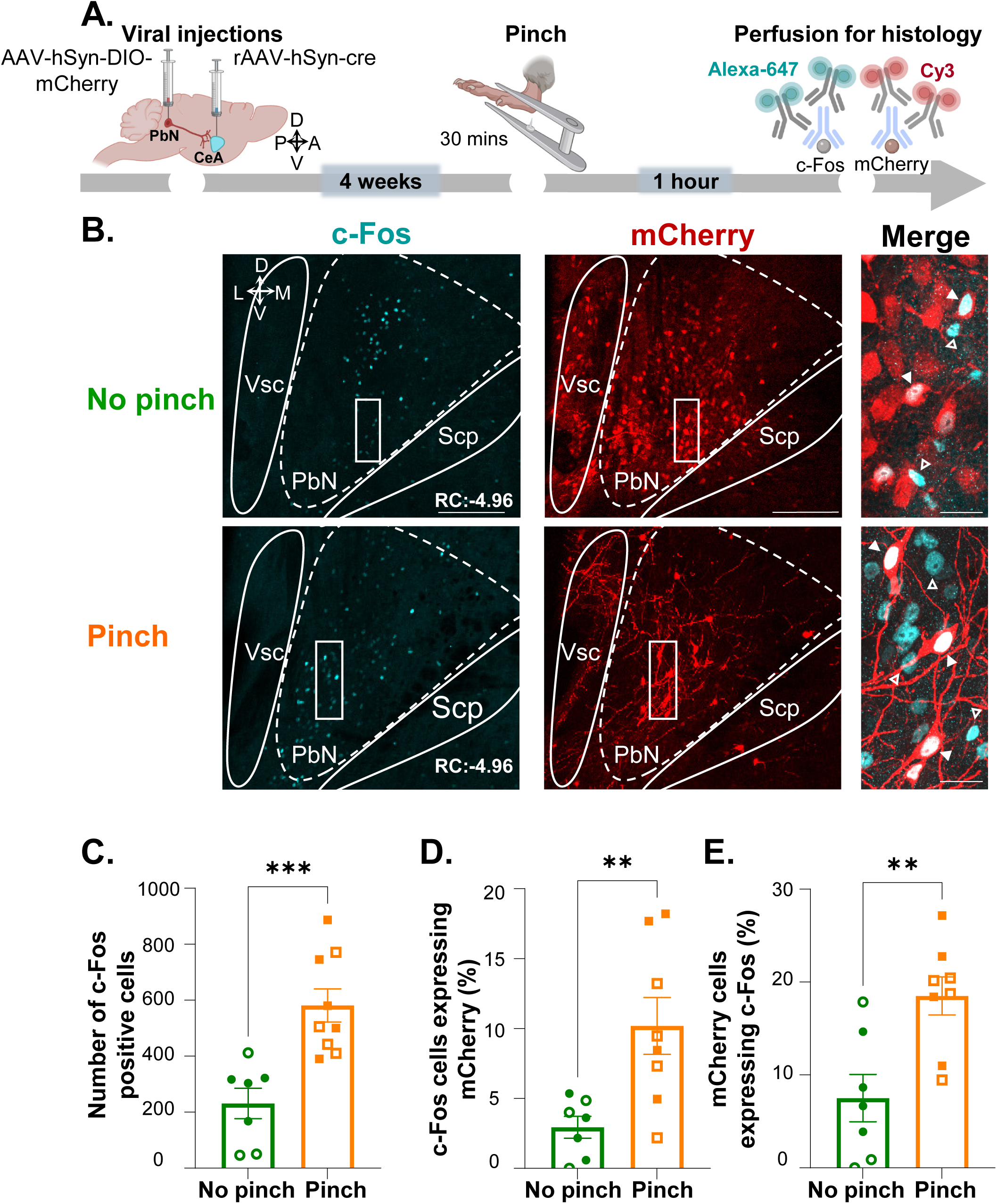
Peripheral noxious stimulation induces c-Fos expression in CeA-projecting PbN neurons. **(A)** Timeline of experiments. Male and female C57BL/6J mice were stereotaxically injected with AAV8-hSyn-DIO-mCherry into the PbN and AAV.hSyn.HI.eGFP-Cre into the CeA. Four weeks after viral injections, we performed pinch on the right hind paw for 30 minutes followed by transcardial perfusion for histology. **(B)** Representative images of PbN slices immunostained for mCherry (11) and c-Fos (cyan). No pinch (top) and pinch (bottom) panels show low magnification images of c-Fos (left) and mCherry (middle). High magnification images of merged signals in area delineated by white box are shown in the right panel. Scale bars represent 200 µm for low magnification and 25 µm for high magnification images. Solid arrows represent cells co-labeled with mCherry and c-Fos while open arrows show c-Fos only positive cells. RC for both animals is -4.96. **(C)** Mean ± SEM of c-Fos+ cells of control (no pinch) vs experimental (pinch) mice in the PbN (n= 7 no pinch mice and n=7 pinch mice; 4 slices per mouse. Unpaired two tailed t test: t=4.223, df=14, ***p= 0.0009 for pinch vs no pinch). **(D)** Mean ± SEM percentage of mCherry cells co-labeled with c-Fos in the PbN of pinch vs no pinch animals (Unpaired two tailed t test: t=3.139, df= 13, **p= 0.0078 for pinch vs no pinch). **(E)** Mean ± SEM percentage of c-Fos+ cells co-labeled with mCherry in the PbN of pinch vs no pinch animals (N=7 mice for no pinch and n=8 mice for pinch; 4 slices per mouse; unpaired two tailed t test: t=3.188, df=13, **p= 0.0049 for pinch vs no pinch). Individual mice are represented by scatter points and open symbols represent female mice.

### Chemogenetic inhibition of CeA-projecting PbN neurons decreases cuff-induced hypersensitivity without affecting baseline nociception

To test for a causal relationship between the activity of CeA-projecting PbN neurons and injury-induced hypersensitivity, we used an intersectional chemogenetic approach where we stereotaxically co-injected a cre-expressing retrograde AAV into the CeA and an AAV encoding the cre-dependent inhibitory designer receptor exclusively activated by designer drugs (DREADD) hM4Di into the PbN (**Figure 4A-E**). To validate our intersectional chemogenetic approach, we prepared acute amygdala slices from mice expressing hM4Di in the PbN and performed whole-cell current-clamp recordings before and after bath-administered clozapine-N-oxide (CNO; 10 μM) or saline control (**Figure 4B**). As expected, bath-administered CNO, but not saline, significantly inhibits neuronal firing in hM4Di-transduced cells (**Figure 4C**), confirming CNO-mediated inhibition of CeA-projecting PbN neurons.

**Figure 4:**
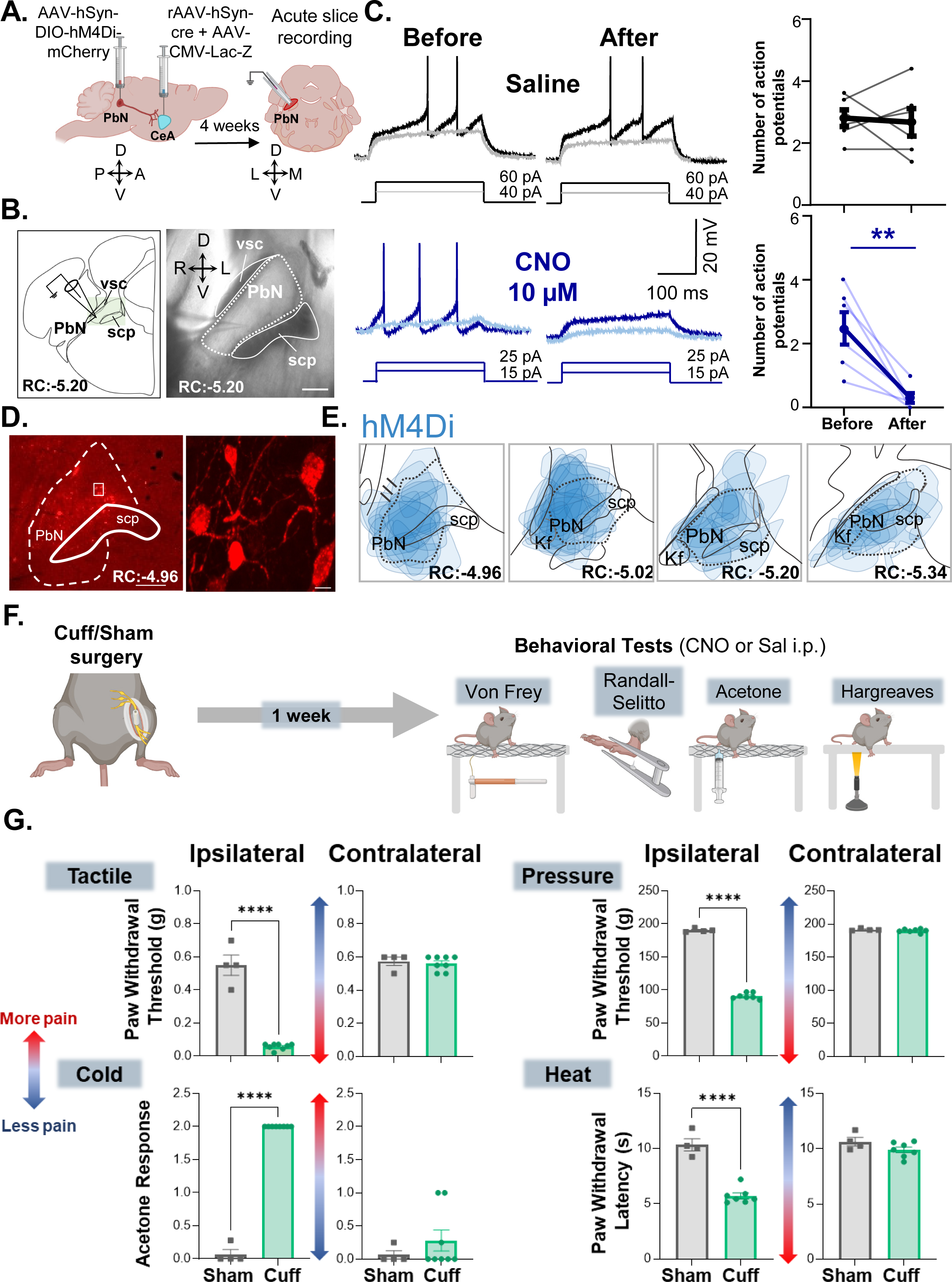
Validation of chemogenetic intersectional approach and behavioral assays. **(A)** Timeline of acute slice electrophysiology experiments. Male C57BL/6J mice were stereotaxically injected with AAV8-hSyn-DIO-hMD4i into the PbN and AAV.hSyn.HI.eGFP-Cre into the CeA. Acute slice recordings were obtained from hMD4i-transduced PbN neurons 4 weeks after the injection. **(B)** Left panel: schematic description (left) and representative differential interference contrast image (9) of coronal brain slice with recording electrode in PbN. Scale bar represents 150 µm. RC: -5.20. **(C)** Representative voltage traces of recordings obtained from hMD4i-transduced PbN neurons in response to depolarizing current injections before (left) and after (9) bath application of saline (top) or 10µM CNO (bottom). Mean ± SEM of the number of action potentials before and after bath applications. (n=6 cells per treatment; paired two tailed T test: t=4.318, df=5 **p=0.0076). **(D)** Representative image of the AAV8-hSyn-DIO-hMD4i injection into the PbN in a coronal brain slice. Immunofluorescence for mCherry is shown in red. Left panel shows a low magnification image and right panel a high magnification of RC: -4.96. Scale bar for left panel is 125 µM and 10 µM for right panel. **(E)** Drawings showing the rostro-caudal distribution of AAV8-hSyn-DIO-hMD4i injection into the PbN. **(F)** Experimental timeline of behavioral experiments. Sciatic nerve sham surgery was performed in male and female C57BL/6J and male Swiss Webster mice. Following 1 week of recovery, Acetone, Hargreaves, Von Frey and Randall-Selitto tests were used to address cold, heat, tactile and pinch stimulation, respectively. **(G)** Mean ± SEM paw withdrawal threshold after tactile or pinch stimulation, acetone response score and withdrawal latency after heat stimulation of the hind paw ipsilateral or contralateral to sham and cuff treatments. Individual mice are represented by scatter points. (n= 4 mice for sham and n=8 mice for cuff in all tests). Unpaired two tailed t test was performed for all behavioral assays. **Tactile**: t= 11.70, df= 10, ****p value<0.0001; **pressure:** t=39.42, df=9, ****p<0.0001; **cold:** t= 41.10, df= 10, ****p<0.0001; **heat**: t=8.536, df=9, ****p <0.0001). Abbreviations: ventral spinocerebellar tract (vsc), superior cerebellar peduncle (scp), Kölliker-Fuse nucleus (kf).

To assess sensitivity to cold, heat, tactile, and pressure stimulation, we used the cuff model of neuropathic pain Benbouzid*, et al.* (40) coupled with acetone, Hargreaves, Von Frey filaments, and Randall Selitto tests, respectively (**Figures 4F**). Control mice received sham sciatic nerve surgeries. Consistent with previous studies (40), sciatic cuff implantation produces robust hypersensitivity in all four modalities tested (**Figure 4G**). Thus, compared to sham mice, cuff mice exhibited significantly (p < 0.0001) lower withdrawal thresholds to tactile and pressure stimulation, higher response scores to cold stimulation, and lower withdrawal latencies to heat stimulation of the hind paw ipsilateral to nerve treatment. Responses to peripheral stimulation of the hind paw contralateral to nerve treatment were indistinguishable between cuff and sham mice demonstrating that hypersensitivity is restricted to the ipsilateral paw.

Using the cuff neuropathic pain model, we next evaluated the effects of chemogenetic inhibition of CeA-projecting PbN neurons on hypersensitivity to tactile, pressure, cold and heat stimulation (**Figure 5A**). Our experiments revealed that cuff-induced hypersensitivity to all modalities is reversed following chemogenetic inhibition of CeA-projecting PbN neurons (**Figure 5B**). Thus, significantly (p < 0.001) higher paw withdrawal thresholds to tactile and pressure stimulation, lower response scores to acetone, and higher withdrawal latencies to heat stimulation were observed after CNO-mediated chemogenetic inhibition of CeA-PbN neurons, compared to before CNO treatment or control saline-treated mice. No measurable effect was seen after CNO treatment in mice stereotaxically injected in the PbN with the control fluorophore mCherry indicating that CNO by itself does not alter the nociceptive behaviors evaluated. Responses in the hind paw contralateral to sciatic cuff implantation were also unaltered by chemogenetic inhibition of the PbN→CeA pathway demonstrating that modulation of behavioral responses to noxious stimuli is restricted to injured states.

**Figure 5:**
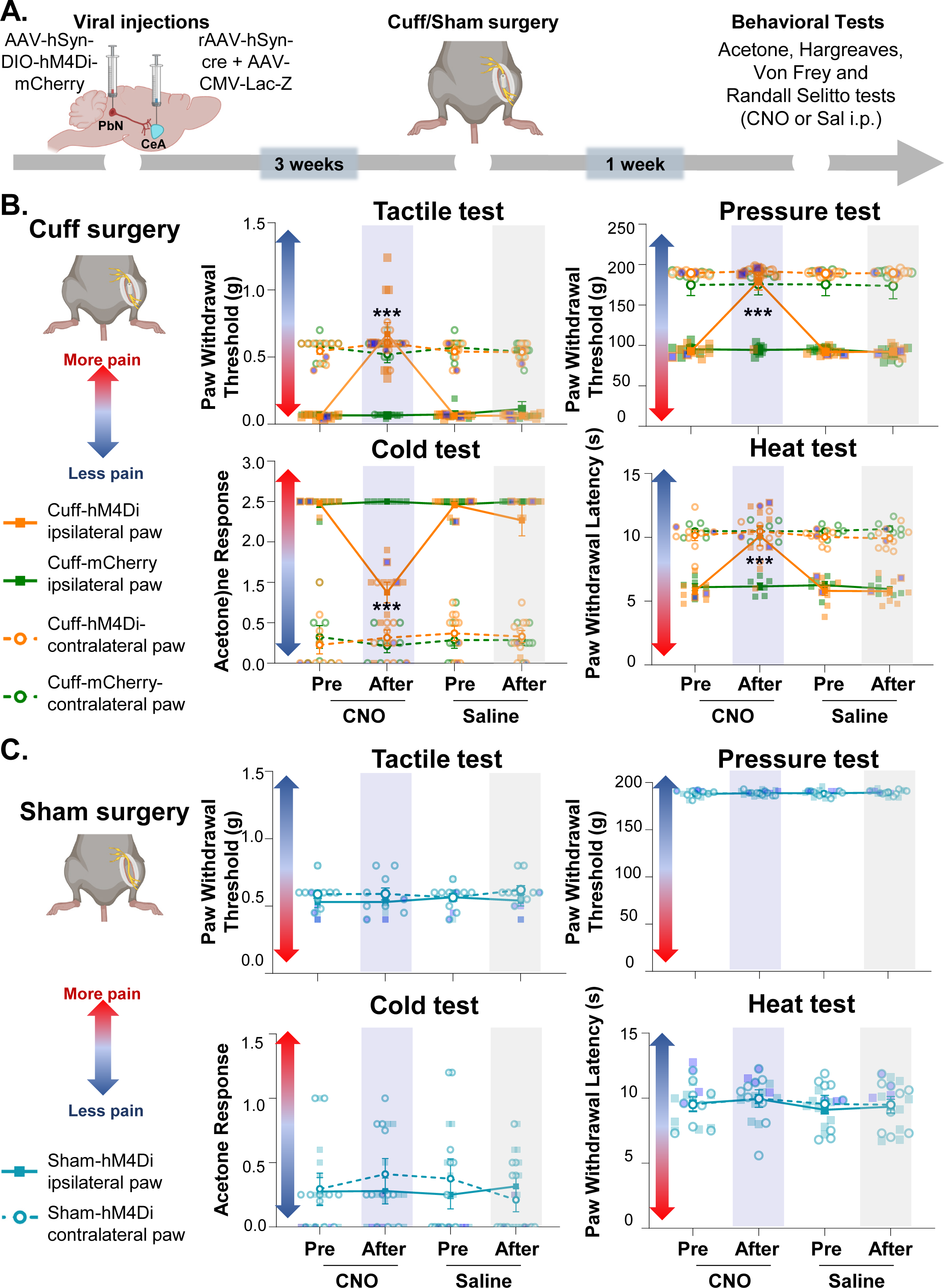
Chemogenetic inhibition of CeA-projecting PbN neurons reverses cuff-induced hypersensitivity without affecting baseline nociception. **(A)**: Experimental timeline Male and female C57BL/6J mice were stereotaxically co-injected with AAV8-hSyn-DIO-hMD4i or AAV8-hSyn-DIO into the right PbN and a mix (1:1) of AAV.hSyn.HI.eGFP-Cre and pAAV.CMV.LacZ.bGH into the right CeA. Sciatic nerve cuff or sham surgery was performed 3 weeks after viral injections. Following 1 week of recovery, Acetone, Hargreaves, Von Frey and Randall-Selitto tests were used to address cold, heat, tactile and pressure stimulation, respectively. Mice were intraperitoneally (i.p.) injected with CNO and saline prior to behavior testing in a counterbalanced way. **(B):** Mean ± SEM paw withdrawal threshold after tactile and pinch stimulation, acetone response score, and withdrawal latency of the hind paw contralateral or ipsilateral to cuff treatment 30 minutes before and after i.p. injection of CNO (blue bar) or Saline (gray bar) in mice injected with the inhibitory DREADD (hM4Di) or the control virus (mCherry) into the PbN and the retrograde tracer into the CeA. Individual mice are represented by scatter points and female mice are identified in purple (n=11 mice for cuff-hM4Di ipsilateral and contralateral paws, n=9 mice for cuff-mCherry ipsilateral and contralateral paws). **(C):** Mean ± SEM paw withdrawal threshold after tactile and pinch stimulation, acetone response score and withdrawal latency after heat stimulation of the hind paw ipsilateral or contralateral to sham treatment before and 30 minutes after i.p. injection of CNO (blue bar) or saline (gray bar) in mice injected with the inhibitory DREADD (hM4Di) into the PbN and the retrograde tracer into the CeA. (n=12 mice for both, ipsilateral and contralateral paws). Two-way repeated measures ANOVA followed by Tukey’s multiple comparison test was performed for all behavioral assays. **Tactile**: brain and paw treatment: F(_3,36_) = 171.3, ****p<0.0001; pre and after i.p. treatment: F_(1.594,57.39)_ = 20.66, ****p<0.0001; interaction: F_(9, 108)_ = 22.33, ****p<0.0001; post hoc: 95.00% CI of diff.= -0.8774 to - 0.3518, ***p=0.0002 for before and after CNO in cuff-hMd4i mice. **Pressure**: brain and paw treatment: F_(3,28)_ = 54.18, ****p<0.0001; pre and after i.p. treatment: F_(1.362,38.13)_ = 4, ****p<0.0001; interaction: F_(9,84)_ = 42.32, ****p<0.0001; post hoc: 95.00% CI of diff.= - 120.4 to -52.46, ***p=0.0002 for before and after CNO in cuff-hMd4i mice. **Cold:** brain and paw treatment: F_(3,30)_ = 352.6, ****p<0.0001; pre and after i.p. treatment: F_(2.734, 82.01)_ = 7.190, ***p=0.0001; interaction: F_(9,90)_ = 7.536, ****p<0.0001; post hoc: 95.00% CI of diff.= 0.6732 to 1.537, ***p=0.0001 for before and after CNO in cuff-hMd4i mice. **Heat:** brain and paw treatment: F_(3, 29)_ = 36.00, ****p<0.0001; pre and after i.p. treatment: F_(1.969, 57.09)_ = 55.02, ***p<0.0001; interaction: F_(9,87)_ = 15.57, ****p<0.0001; post hoc: 95.00% CI of diff.= -6.370 to -2.230, ***p=0.0007 for before and after CNO in cuff-hMd4i mice.

In contrast, and consistent with previous reports (12, 31, 41), evaluation of control (sham) mice showed that responses to peripheral stimulation were comparable to before and after chemogenetic inhibition of CeA-projecting PbN neurons independently of the modality or the hind paw tested (**Figure 5C**). Together, these results demonstrate that activity of CeA-projecting PbN neurons is necessary for hypersensitivity in a model of neuropathic pain but does not modulate baseline responses to cold, heat, tactile and pressure stimulation in uninjured states.

### Chemogenetic inhibition of CeA-projecting PbN neurons reduces licking behavior after intraplanar formalin injection

To assess the function of the PbN→CeA pathway in a context of inflammatory pain, we measured spontaneous nociceptive responses to 2-3% formalin injection into the hind paw of mice following inhibition of CeA-projecting PbN neurons using the intersectional chemogenetic strategy described in the section above (**Figure 6A**). Evaluation of the time spent in nociceptive behaviors as a function of time after formalin injection showed a stereotypical biphasic response to formalin in all animals tested (42) (**Figure 6B**). The time spent in nociceptive behaviors during the second phase, however, was significantly (p < 0.05) lower after CNO-mediated chemogenetic inhibition of CeA-projecting PbN neurons than in control saline-treated mice that also received stereotaxic injection of an AAV encoding the hM4Di inhibitory DREADD (**Figure 6B-D**).

**Figure 6:**
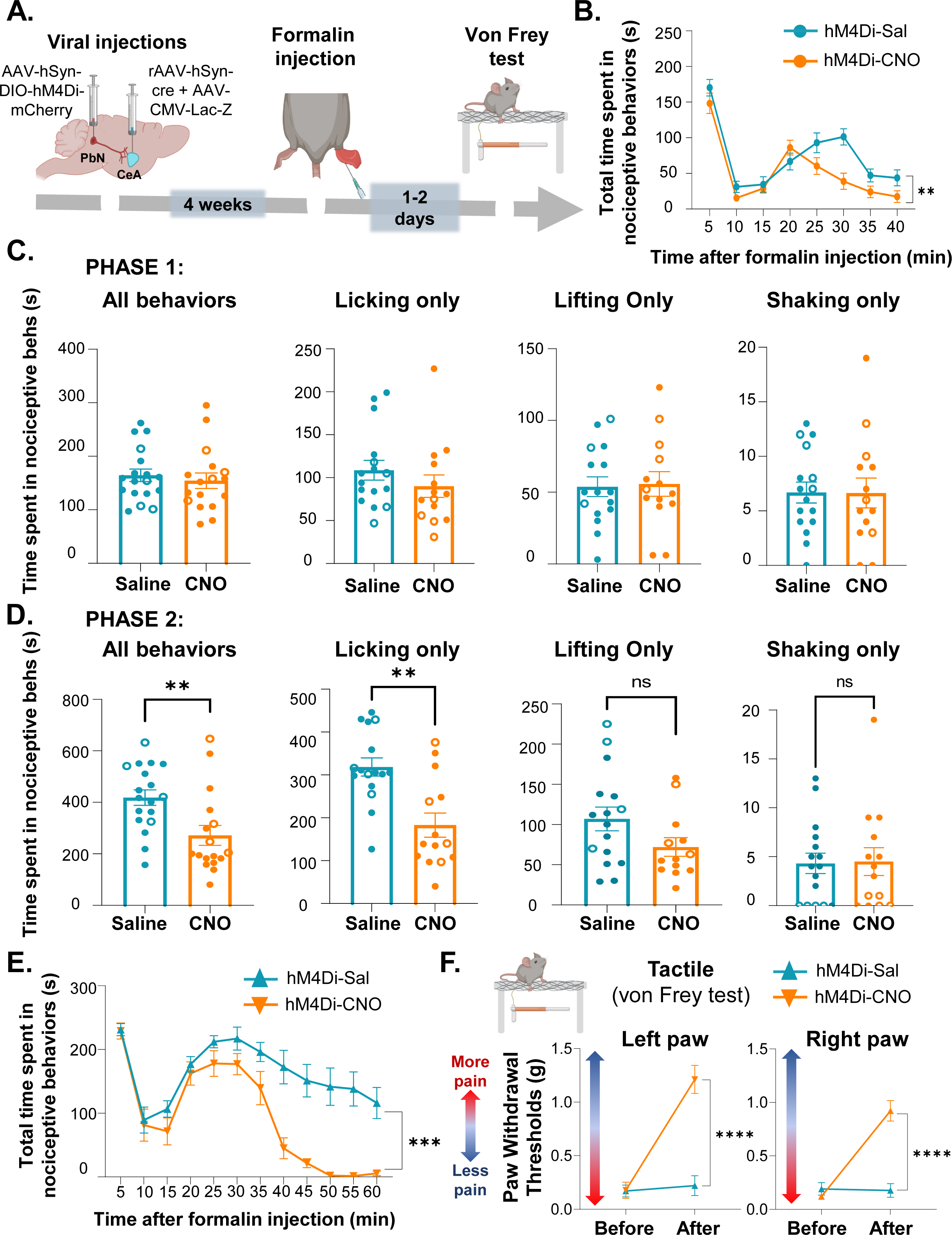
Inhibition of CeA-projecting PbN neurons reduces licking behaviors after intraplantar formalin injection. **(A)**: Experimental timeline. Male and female C57BL/6J and male Swiss Webster mice were stereotaxically co-injected with AAV8-hSyn-DIO-hMD4i-mCherry into the right PbN and a mix (1:1) of AAV.hSyn.HI.eGFP-Cre and pAAV.CMV.LacZ.bGH into the right CeA. 4 weeks after viral injections, mice were injected in the hind paw with 2-3% formalin 30 minutes after CNO or saline i.p. injections. Licking, lifting, and shaking nociceptive behaviors were measured for 40 mins. **(B):** Mean ± SEM of time spent in nociceptive behaviors as a function of time after formalin injection in C57BL/6J mice. Two-way repeated measures ANOVA was performed; (i.p. treatment: F_(1, 33)_ = 9.001, **p=0.0051; time after formalin injection: F_(4.495, 148.3)_ = 41.25, ****p<0.0001; interaction: F_(7,231)_ = 2.779, **p=0.0086). **(C-D):** Mean ± SEM of time spent in distinct nociceptive behaviors during phase 1 (C) and phase 2 (D). Phase 1 is defined as 0-5 mins post formalin injection and phase 2 as 5-40 mins post formalin injection. Individual mice are represented by scatter points and open symbols represent female mice (n=18 mice for saline and n=17 mice for CNO for all experiments). Unpaired two tailed t test was performed. Sum of all nociceptive behaviors: t= 3.0287, df= 33, **p= 0.0048. Licking behavior only: t= 3.891, df= 28, ***p= 0.0006. **(E):** Mean ± SEM of time spent in nociceptive behaviors as a function of time after formalin injection in Swiss Webster mice. Two-way repeated measures ANOVA was performed; (i.p. treatment: F_(1, 19)_ =16.42, ***p=0.0007; time after formalin injection: F_(4.478, 85.09)_ = 34.56, ****p<0.0001; interaction: F_(11,209)_ = 7.186, ****p<0.0001). **(F):** Mean ± SEM paw withdrawal threshold after tactile stimulation before and after CNO or Saline in both paws 1 day after formalin test. Two-way repeated measures ANOVA followed by Sidaks multiple comparisons test was performed. **Left paw:** i.p. treatment: F_(1, 11)_ =19.09, **p=0.0011; before and after i.p.: F_(1, 11)_ = 85.77, ****p<0.0001; interaction: F_(1,11)_ = 70.68, ****p<0.0001; post hoc: 95.00% CI of diff.= -1.257 to -0.8122, ****p<0.0001 for before and after CNO in hMd4i mice. **Right paw:** i.p. treatment: F_(1, 11)_ =17.48, **p=0.0015; before and after i.p.: F_(1, 11)_ = 78.68, ****p<0.0001; interaction: F_(1,11)_ = 84.51, ****p<0.0001; post hoc: 95.00% CI of diff.= - 0.9699 to -0.6331, ****p<0.0001 for before and after CNO in hMd4i mice.

Nociceptive responses to formalin are defined as lifting, licking, or shaking of the injected hind paw (43, 44). Recent studies suggested that modulation of distinct behavioral responses to noxious stimuli are circuit specific (45–47). To determine if modulation of spontaneous nociceptive responses to formalin by CeA-projecting PbN neurons is behavior-specific, we analyzed the time spent licking, lifting, or shaking separately in both phase 1 and phase 2 of the formalin test. Our results showed that all nociceptive responses during phase 1 of the formalin test are comparable in control saline-treated mice and in mice with CNO-mediated chemogenetic inhibition of CeA-projecting PbN neurons (**Figure 6C**). In contrast, during phase two, inhibition of CeA-projecting PbN neurons significantly (p < 0.01) decreased licking behaviors without measurably affecting lifting and shaking behaviors (**Figure 6D**). Swiss Webster mice showed similar decreases in nociceptive responses to 5% hind paw formalin injection following chemogenetic inhibition of CeA-projecting PbN neurons (**Figure 6E**). Additionally, CNO-mediated inhibition of these cells significantly (p <0.0001) decreased formalin-induced hypersensitivity to tactile stimulation (**Figure 6F**). Thus, CNO-treated mice displayed higher paw withdrawal thresholds compared to saline-treated control mice. Altogether, our results demonstrate that activity of CeA-projecting PbN neurons is necessary for formalin-induced licking behavioral responses and tactile hypersensitivity but that lifting and shaking responses to formalin are not dependent on the PbN→CeA pathway.

### Chemogenetic activation of CeA-projecting PbN neurons results in bilateral hypersensitivity in a modality specific way

To test whether activation of CeA-projecting PbN neurons is sufficient to induce hypersensitivity to tactile, pressure, cold, and heat stimulation, we used an intersectional chemogenetic approach in which we co-injected a cre-expressing retrograde AAV into the CeA and an AAV encoding the cre-dependent excitatory DREADD hM3Dq into the PbN (**Figure 7A**). As illustrated in **Figure 7B**, the number of transduced CeA-projecting PbN neurons co-expressing the neuronal activity marker c-Fos is significantly (p < 0.01) higher in brain slices from mice following CNO-mediated activation of the hM3Dq DREADD, compared to the number of c-Fos+ cells in slices from control saline-treated mice. These results confirmed we can selectively activate CeA-projecting PbN neurons using this intersectional chemogenetic strategy.

**Figure 7:**
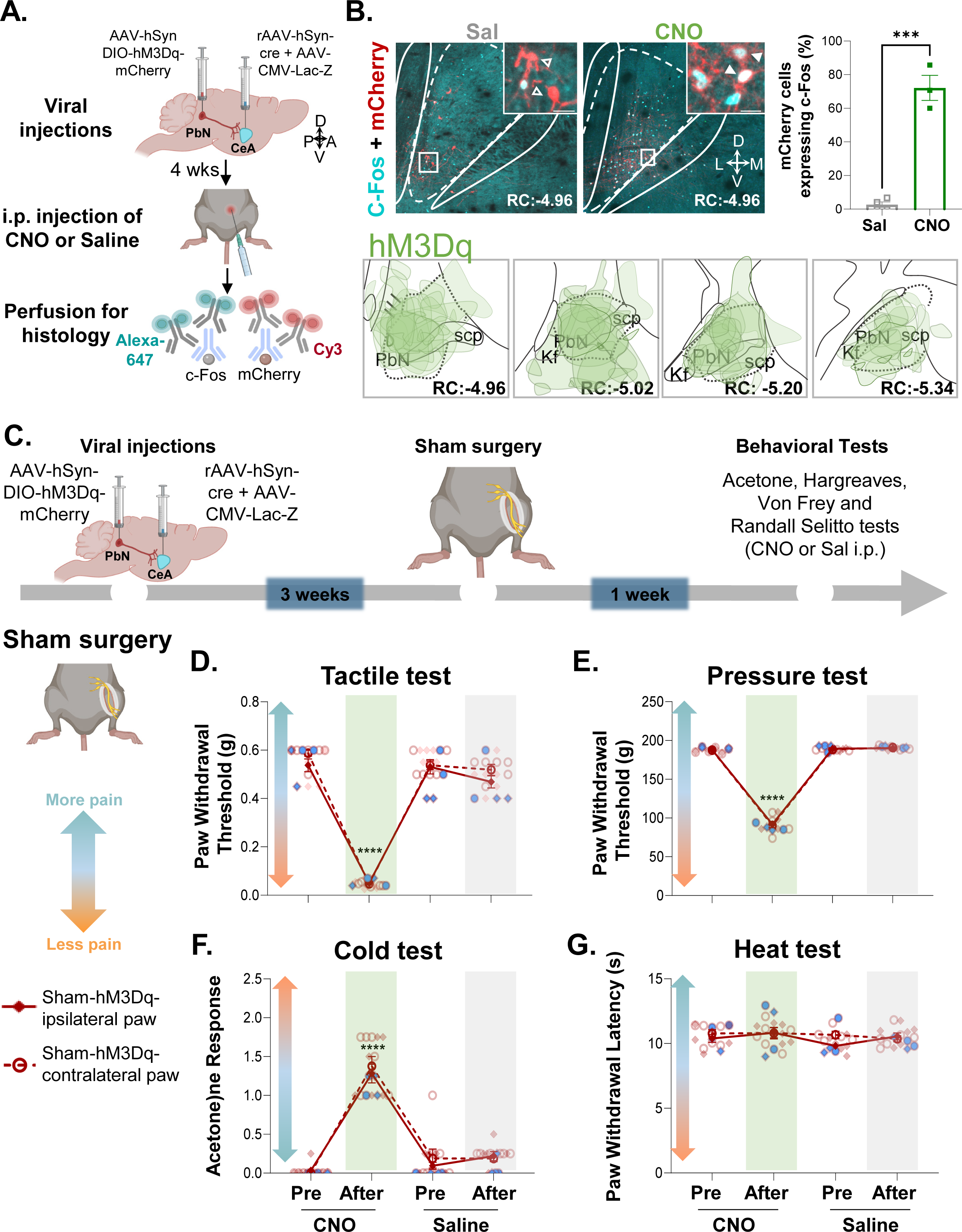
Chemogenetic activation of CeA-projecting PbN neurons induces bilateral hypersensitivity in uninjured mice in a modality specific manner. **(A)**: Timeline of chemogenetic activation validation. Male C57BL/6J mice were stereotaxically injected with AAV8-hSyn-DIO-hMD3q into the right PbN and a mix (1:1) of AAV.hSyn.HI.eGFP-Cre and pAAV.CMV.LacZ.bGH into the right CeA. 4 weeks after, mice were intraperitoneally injected with either CNO or saline and perfused 30-45mins after for immunohistology purposes. **(B):** Representative images of PbN slices of stereotaxically injected mice after saline (left) and CNO (9) i.p. injections. Immunofluorescence for mCherry (representing hM3Dq-transduced cells) is shown in red and for c-Fos in cyan color. Insets: solid arrows represent mCherry+ cells colocalized with c-Fos while open arrows represent mCherry+ only cells. Scale bar represents 150 µm for low magnification and 10 µm for insets. RC for both pictures: -4.96. Right panel shows mean ± SEM percentage of mCherry+ cells expressing c-Fos (n=4 mice for saline, n=3 mice for CNO; 4 slices per mouse). Unpaired two tailed t test was performed. T= 10.64, df= 5, ***p=0.0001. Bottom panel shows drawings of the rostro-caudal distribution of the AAV8-hSyn-DIO-hMD3q injection into the PbN. **(C):** Experimental timeline of behavioral experiments. Male and female C57BL/6J mice were stereotaxically injected with AAV8-hSyn-DIO-hM3Dq into the right PbN and rAAV-hSyn-cre into the right CeA. Sciatic nerve sham surgery was performed 3 weeks after viral injections. Following 1 week of recovery, Acetone, Hargreaves, Von Frey and Randall-Selitto tests were used to address cold, heat, tactile and pinch stimulation, respectively. Mice were intraperitoneally (i.p.) injected with CNO and saline prior to behavior testing in a counterbalanced way**. (D-G):** Mean ± SEM paw withdrawal threshold after tactile and pinch stimulation, acetone response score and withdrawal latency after heat stimulation of the hind paw ipsilateral or contralateral to sham treatment before and 30 minutes after i.p. injection of CNO (green bar) or saline (gray bar) in mice injected with the excitatory DREADD (hM3Dq) into the PbN and the retrograde tracer into the CeA. Blue symbols represent female mice and individual mice are represented by scatter points (n= 8 mice per treatment for all tests). Two-way repeated measures ANOVA followed by Tukey’s multiple comparison test was performed for all behavioral assays. **Tactile**: contra/ipsi paw: F_(1,14)_ = 2.979, p=0.1068; i.p. treatment: F_(2.487,34.82)_ = 296, ****p<0.0001; interaction: F_(3,42)_ = 1.002, p=0.4012; post hoc: 95.00% CI of diff.= 0.3999 to 0.5751, ****p<0.0001 for before and after CNO in sham-hM3dq mice ipsilateral paw and 95.00% CI of diff.= 0.4969 to 0.5843, ****p<0.0001 for before and after CNO in sham-hM3dq mice contralateral paw. **Pressure**: contra/ipsi paw: F_(1,10)_ = 0.08549, p=0.7760; pre and after i.p. treatment: F_(1.232,12.32)_ = 891.8, ****p<0.0001; interaction: F_(3,30)_ = 0.09099, p=0.96; post hoc: 95.00% CI of diff.= 82.64 to 110.0, ****p<0.0001 for before and after CNO in sham-hM3dq mice ipsilateral paw and 95.00% CI of diff.= 83.28 to 111.4, ****p<0.0001 for before and after CNO in sham-hM3dq mice contralateral paw. **Cold:** contra/ipsi paw: F_(1,14)_ = 0.1707, p=0.6857; pre and after i.p. treatment: F_(2.209, 30.93)_ = 147.8, ****p<0.0001; interaction: F_(3, 42)_ = 0.5185, p=0.6718; post hoc: 95.00% CI of diff.= -1.600 to -0.9003, ****p<0.0001 for before and after CNO in sham-hM3dq mice ipsilateral paw and 95.00% CI of diff.= -1.789 to -0.9612, ****p<0.0001 for before and after CNO in sham-hM3dq mice contralateral paw.

We then evaluated the effects of chemogenetic activation of CeA-projecting PbN neurons on sensitivity to tactile, pressure, cold, and heat stimulation of the hind paw using the Von Frey, Randall Selitto, acetone, and Hargreaves tests, respectively (**Figure 7C**). Our results revealed that CNO-mediated activation of CeA-projecting PbN neurons is sufficient to induce bilateral tactile, pressure, and cold hypersensitivity in uninjured mice (**Figure 7D-F**). Thus, significantly (p < 0.01) lower paw withdrawal thresholds to tactile and pressure stimulation and higher acetone response scores were observed following CNO treatment, compared to pre-treatment responses and responses in control saline-treated mice. In marked contrast, however, paw withdrawal latencies in response to heat stimulation were unaltered by CNO-mediated activation of CeA-projecting PbN neurons (**Figure 7G**).

Similar results were observed in the paw contralateral to sciatic nerve cuff implantation, with CNO-treated cuff mice displaying significant (p < 0.01) hypersensitivity to tactile, pressure, and cold, but not heat stimulation, compared to pretreatment responses and responses in control saline-treatment (**Figure 8**). No measurable effects were observed in the paw ipsilateral to the cuff implantation after CNO treatment, possibly due to a hypersensitivity ceiling effect. Collectively, our results demonstrate that CeA-projecting PbN neurons are sufficient to induce bilateral hypersensitivity in the absence of injury in a modality-specific way.

**Figure 8:**
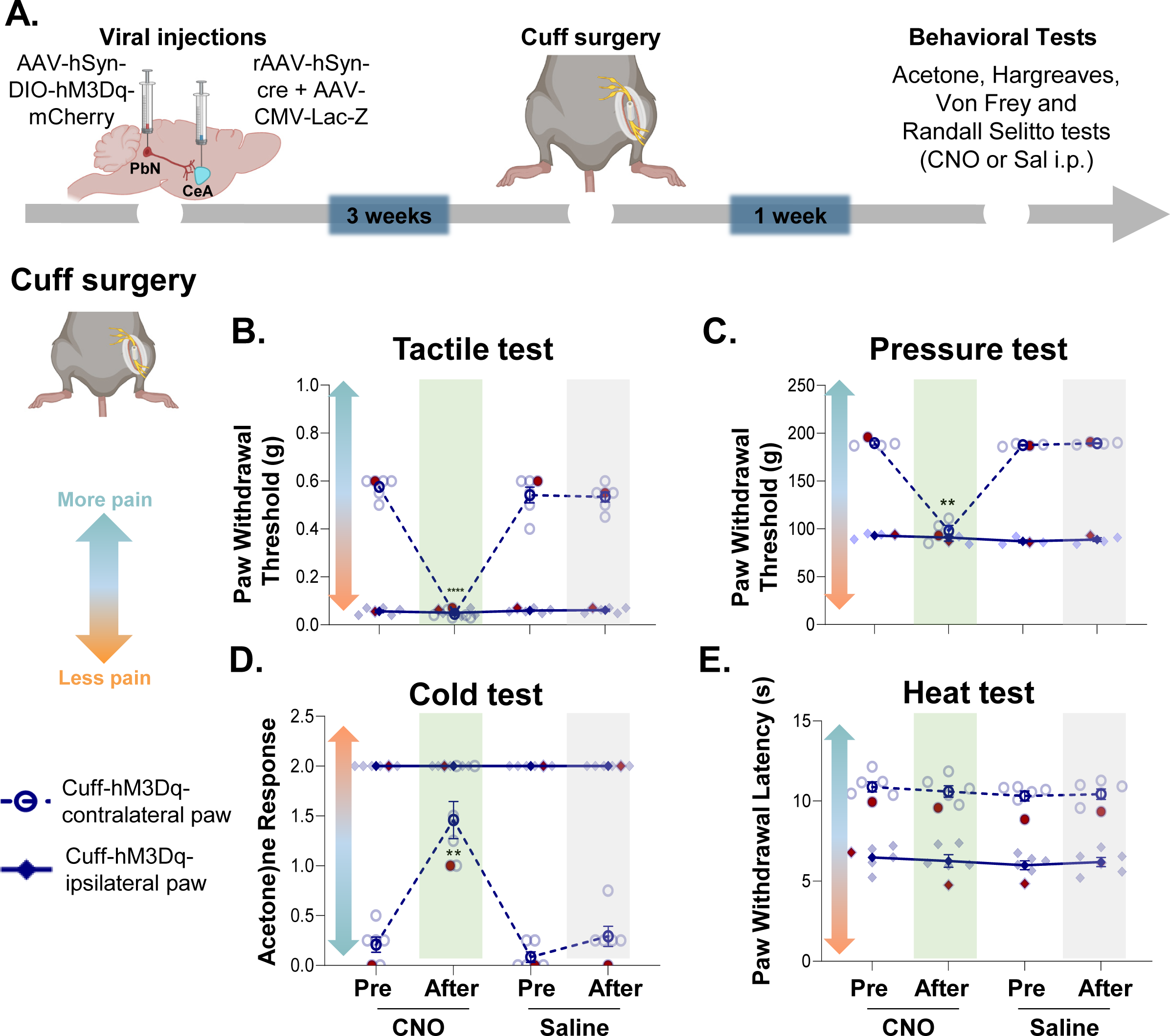
Chemogenetic activation of CeA-projecting PbN neurons induces contralateral hypersensitivity in a modality specific manner in injured mice. **(A)**: Experimental timeline. Male and female C57BL/6J mice were stereotaxically injected with AAV8-hSyn-DIO-hM3Dq into the right PbN and a mix (1:1) of AAV.hSyn.HI.eGFP-Cre and pAAV.CMV.LacZ.bGH into the right CeA. Sciatic nerve cuff surgery was performed 3 weeks after viral injections. Following 1 week of recovery, Acetone, Hargreaves, Von Frey and Randall-Selitto tests were used to address cold, heat, tactile and pinch stimulation, respectively. Mice were intraperitoneally (i.p.) injected with CNO and saline prior to behavior testing in a counterbalanced way. **(B-E):** Mean ± SEM paw withdrawal threshold after tactile and pinch stimulation, acetone response score and withdrawal latency after heat stimulation of the hind paw ipsilateral or contralateral to cuff treatment before and 30 minutes after i.p. injection of CNO (green bar) or saline (gray bar) in mice injected with the excitatory DREADD (hM3Dq) into the PbN and the retrograde tracer into the CeA. Red symbols represent female mice and individual mice are represented by scatter points (n= 8 mice per treatment for all tests). Two-way repeated measures ANOVA followed by Tukey’s multiple comparison test was performed for all behavioral assays. **Tactile**: ipsi/contra paw: F_(1,10)_ = 842.2, ****p<0.0001; pre and after i.p. treatment: F_(2.236,22.36)_ =151.5, ****p<0.0001; interaction: F_(3,30)_ = 141.8, ****p<0.0001; post hoc: 95.00% CI of diff.= 0.4526 to 0.6066, ****p<0.0001 for before and after CNO in cuff-hM3dq contralateral paw mice. **Pressure**: ipsi/contra paw: F_(1,6)_ = 1610, ****p<0.0001; pre and after i.p. treatment: F_(1.525,9.149)_ = 131.3, ****p<0.0001; interaction: F_(3,18)_ = 139.2, ****p<0.0001; post hoc: 95.00% CI of diff.= 61.21 to 122.3, **p=0.0022 for before and after CNO in cuff-hM3dq contralateral paw mice. **Cold:** ipsi/contra paw: F_(1,10)_ = 289.6, ****p<0.0001; pre and after i.p. treatment: F_(1.868, 18.68)_ = 52.64, ****p<0.0001; interaction: F_(3,30)_ = 52.64, ****p<0.0001; post hoc: 95.00% CI of diff.= -1.783 to -0.7174, **p=0.0013 for before and after CNO in cuff-hM3dq contralateral paw mice.

## Discussion

The PbN→CeA is one of the major ascending pain pathways characterized (5–7, 30). Previous studies have shown that manipulations of PbN neurons in this pathway modulate affective-motivational aspects of pain but not baseline somatosensory responses to peripheral noxious stimulation (12, 31, 32, 36, 41). The functionality of this pathway in injury-induced pain sensitization, however, remains unknown. In the present study, we addressed this gap in knowledge using optogenetic assisted circuit mapping in *ex-vivo* brain slices coupled with *in-vivo* behavioral pain assays, intersectional chemogenetic approaches, and histology in two mouse models of persistent pain. We show that CeA-projecting PbN neurons are activated after peripheral noxious stimulation and activity of this pathway is necessary for injury-induced behavioral hypersensitivity, but not baseline nociception (**Figure 9**). We further show that chemogenetic activation of this pathway is sufficient to drive hypersensitivity in the absence of injury. These results are consistent with the growing body of evidence supporting the function of the PbN→CeA pathway in pain processing and further demonstrate that plasticity in this pathway contributes to the amplification of pain-like responses in pathological pain states.

**Figure 9:**
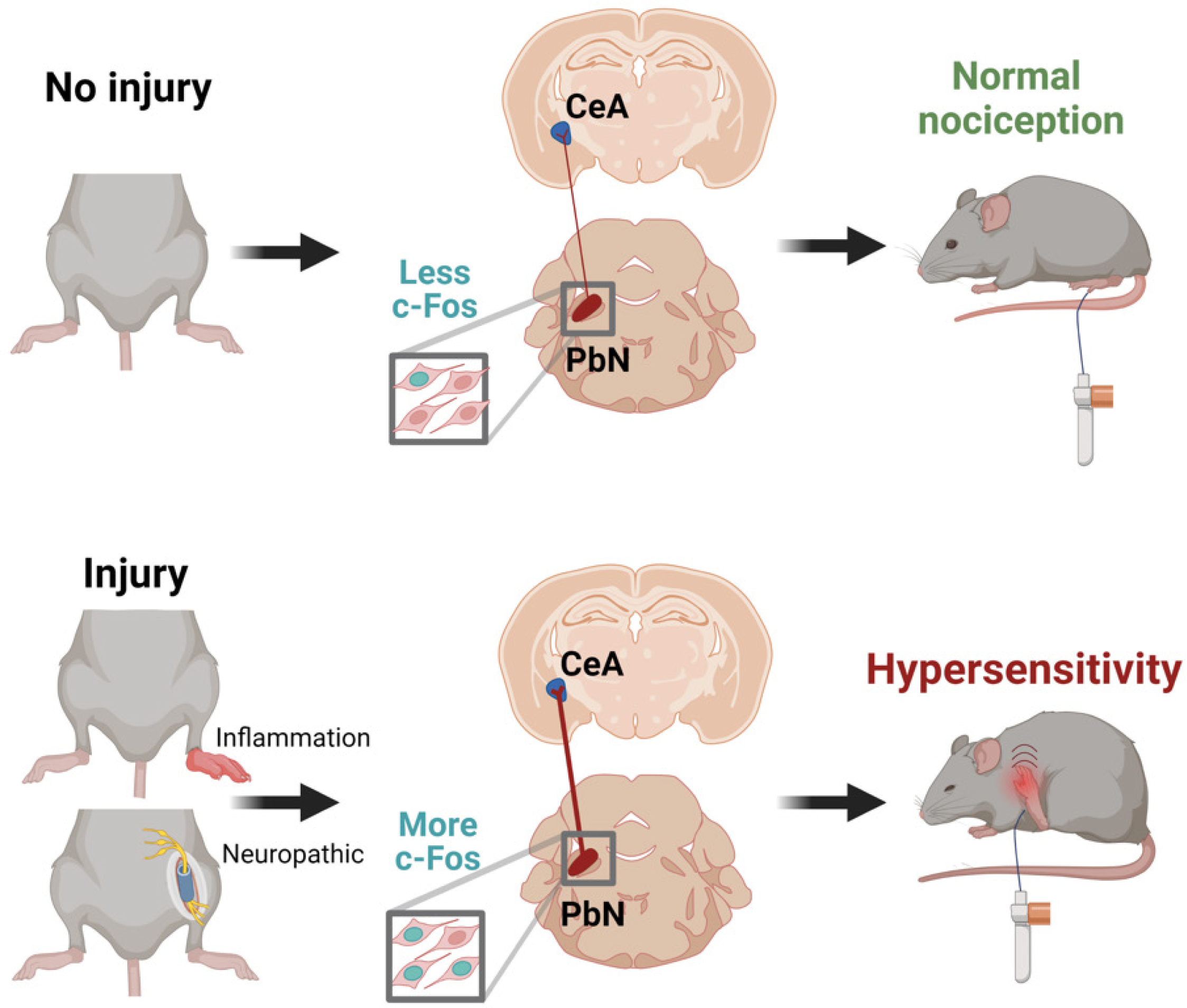
Summary of results and proposed model. Activity of CeA-projecting PbN neurons does not contribute to baseline nociception in uninjured states whereas increases in neuronal activity in this pathway drive peripheral hypersensitivity in injured states.

### Activation of CeA-projecting PbN neurons by peripheral noxious stimulation

Our histological experiments show a subpopulation of noxious-activated PbN neurons project to the CeA (**Figure 3D-E**). These results were initially surprising given previous reports demonstrating spinal nociceptive inputs do not target PbN neurons that project to the CeA (36, 48, 49). Classical studies have shown, however, that injection of a transneuronal retrograde tracer into the CeA results in progressive infection of a multi-synaptic circuit that includes neurons in the parabrachial nucleus, medulla, and dorsal horn (7). These results demonstrate the dorsal horn of the spinal cord is in fact anatomically linked to PbN neurons that project to the CeA. Consistently, recent *ex-vivo* electrophysiological experiments in acute brain slices revealed the existence of a functional excitatory microcircuit within the PbN that connects dynorphin-expressing neurons that receive spinal nociceptive inputs with neurons that project to the CeA (12). The results from our c-Fos experiments showing activation of the PbN→CeA pathway by peripheral noxious stimulation, coupled with the anatomical and electrophysiological studies described above, suggest that pain-induced activation of the PbN→CeA pathway occurs via polysynaptic excitatory inputs from the dorsal horn of the spinal cord to PbN neurons that project to the CeA.

### CeA-projecting PbN neurons as major contributors to injury-induced pain sensitization but not baseline nociception

*In-vivo* and *ex-vivo* electrophysiological studies have shown injury-induced sensitization and potentiation of synaptic transmission in the PbN→CeA pathway in several models of persistent pain, suggesting that activity of this pathway contributes to pain-related behavioral responses (4, 20, 21, 26, 27). Our findings demonstrating that chemogenetic inhibition of CeA-projecting PbN neurons reverses injury-induced hypersensitivity (**Figure 5B**) and chemogenetic activation drives hypersensitivity in the absence of injury (**Figure 7**) in both male and female mice provide a causal link for the function of this pathway in pain processing following peripheral inflammation or nerve injury.

The analgesic effects we see following chemogenetic inhibition of the PbN→CeA pathway are specific to injured states. Thus, chemogenetic inhibition of CeA-projecting PbN neurons does not affect withdrawal responses in sham-treated animals (**Figure 5B**) or in the paw contralateral to nerve injury (**Figure 5C**). These results are consistent with previous studies in naïve animals that report no measurable effects on reflexive withdrawal responses to somatosensory stimuli after manipulations of PbN neurons that project to the CeA and express CGRP, tachykinin receptor 1, or mu-opioid receptor (12, 31, 36, 41). The lack of modulation of reflexive responses to somatosensory stimuli by the PbN→CeA pathway in naïve animals, coupled with the reported function of this pathway in the modulation of aversion, threat memory, and pain-related affective-motivational responses (12, 31, 32) led to the conclusion that CeA-projecting PbN neurons contribute to the affective-motivational but not reflexive-defensive somatosensory component of pain. Our experiments demonstrate CeA-projecting PbN neurons indeed contribute to reflexive somatosensory responses but only in the context of injury. These results underscore the importance of including uninjured and injured states in studies addressing mechanisms for pain processing and modulation.

### Modality-specific effects following activation of the PbN→CeA pathway

We show that chemogenetic inhibition of the PbN→CeA pathway reversed injury-induced hypersensitivity to tactile, pressure, cold, and heat stimuli (**Figure 5B**). In contrast, chemogenetic activation of this pathway elicited hypersensitivity to tactile, pressure and cold (**Figure 7D-F**) but not heat stimulation (**Figure 7G**). Modality-specific effects are often observed in studies that manipulate pain circuits (22, 23, 50–52). In addition, lamina I spinoparabrachial neurons have been shown to display modality-selective responses in an *ex-vivo* semi-intact somatosensory preparation (53), further supporting the idea that distinct mechanisms underlie modulation of specific sensory modalities. The mechanisms driving modulation of distinct modalities remain unclear.

### The PbN→CeA pathway contributes to both reflexive-defensive reactions and affective-motivational responses to painful stimuli

Several pain-related responses to cutaneous noxious stimulation have been previously described (45–47, 54). These behavioral responses have been further categorized as reflexive-defensive reactions (paw withdrawal) or affective-motivational responses (licking, biting, extended lifting, or guarding of the stimulated paw and escape responses such as jumping, hyperlocomotion or rearing). A notable discovery from these prior studies is that distinct anatomical circuits modulate reflexive-defensive reactions vs affective-motivational responses to peripheral noxious stimuli (45, 54). Our findings show that inhibition of the PbN→CeA pathway affects licking responses to formalin (**Figure 6**) suggesting this pathway modulates affective-motivational responses to peripheral inflammation. These findings, combined with our results that chemogenetic manipulations of CeA-projecting PbN neurons also affects reflexive paw withdrawal responses following inflammation or nerve injury (**Figures 5-6**), suggest that the PbN→CeA pathway contributes to both reflexive-defensive reactions (paw withdrawal) and affective-motivational responses (licking) in the context of injury.

### Protective and maladaptive functions of the PbN→CeA in response to injury

The PbN has been described as a general alarm system to potential threats that orchestrates behavioral and physiological responses essential for survival (55). Consistent with this, studies have shown PbN neurons are activated by a variety of danger signals and this activation contributes to behavioral responses to threat in a cell-type and circuit-specific manner (14, 31, 56–64). PbN neurons that express Calcitonin gene-related peptide (CGRP) and project to the CeA, for example, have been shown to modulate escape responses to noxious heat and contribute to threat memory (31). Separate studies have also shown that manipulations of CeA-projecting PbN neurons modulate aversion and affective motivational aspects of pain (12, 31, 36, 65, 66).

Maladaptive plasticity in the PbN→CeA pathway has also been proposed to contribute to persistent hypersensitivity in pathological conditions (18, 26), no longer serving a protective function. Consistent with this idea, recent work demonstrated that CGRP signaling in the right PbN→CeA pathway contributes to bladder pain-like behaviors but not pain-related aversion in a mouse model of cystitis (67). The results in the present study expand on these previous findings to show that the PbN→CeA pathway is also a major contributor of hypersensitivity to peripheral stimulation in response to inflammation or nerve injury. Based on these combined findings, we propose that this circuit functions as a biological alarm system that protects against further injury and promotes healing under physiological conditions. In pathological conditions, however, maladaptive changes in the PbN-CeA pathway might contribute to persistent hypersensitivity that is no longer protective.

### Materials and Methods Subjects

Experiments were approved by the Animal Care and Use Committee of the National Institute of Neurological Disorders and Stroke and the National Institute on Deafness and other Communication Disorders with the guidance from the National Institutes of Health (NIH). Adult C57BL/6J or Swiss Webster mice between the age of 8 to 17 weeks old, bred in house or purchased from Jackson Laboratory, were used for all behavioral and histological experiments. The sex of the mice used for each experiment is described in the sections below. The following mouse lines were used for electrophysiology experiments: heterozygous *Prkcd*-cre mice (GENSAT – founder line 011559-UCD) or heterozygous male *Sst*-cre (Jackson Laboratory – founder line 018973) crossed with homozygous Ai9 mice (Jackson Laboratories). Offspring mice were genotyped for cre-recombinase using tail biopsies and PCR (Transnetyx) with the following primers: TTAATCCATATTGGCAGAACGAAAACG (forward) and CAGGCTAAGTGCCTTCTCTACA (reverse). The expression and fidelity of Cre in Som+ and PKCδ+ neurons have been previously described (23, 68). Mice were initially group housed (up to 5 mice per cage) with littermates of the same sex in a vivarium with controlled humidity and temperature under reversed 12 h light/dark cycle (9 pm to 9 am light). Following surgery, pairs of littermate mice of the same sex and pain treatment were transferred to new home cages with perforated Plexiglass dividers to keep one mouse per side. All behavioral tests were performed under red light during the dark period, between the hours of 10 am and 6 pm. Mice received one handling session per day for at least 5 days before the start of behavioral and electrophysiological experiments as previously described (69). During each handling session, mice were allowed to move freely on the hands of the experimenter for approximately 5-8 mins and were then injected with 50-100 µl saline intraperitoneally (i.p.). Animals were randomly assigned to experimental groups and all experiments and analyses were performed blind to experimental treatment.

### Stereotaxic injections

Acute microinjections were performed using a small animal stereotaxic instrument (David Kopf Instruments). Male and female C57BL/6J and Swiss Webster mice were initially anesthetized with 5% isoflurane in preparation for the stereotaxic surgery. After induction, mice were maintained with 2% isoflurane at a flow rate of 0.5 L/min for the duration of surgery. A hand warmer (Hot Rods Hand Warmers) was used for thermal maintenance during the procedure. Stereotaxic injections were performed using 0.5 µl Hamilton Neuros 32-gauge syringes (Neuro model 7000.5 KH) at a flow rate of 0.1 µl/min. The syringe was left in place for an additional 15 minutes to allow for diffusion of virus and to prevent backflow.

Based on previous literature demonstrating hemispheric lateralization of CeA function in the modulation of hypersensitivity, (70–72) stereotaxic injections were performed in the right hemisphere in all experiments. For intersectional chemogenetic experiments, the CeA of C57BL/6J mice was injected with either 0.05 µl of pENN.AAV.hSyn.HI.eGFP-Cre.WPRE.SV40 (Addgene 105540-AAVrg) or 0.07 µl of a 1:1 mixture of pENN.AAV.hSyn.HI.eGFP-Cre.WPRE.SV40 (Addgene 105540-AAVrg) and pAAV.CMV.LacZ.bGH (Addgene 105531-AAV8). During the same surgery, the PbN was injected with 0.1 µl of AAV8-hSyn-DIO-hM4D(Gi)-mCherry (Addgene 44362-AAV8) or AAV8-hSyn-DIO-mcherry (Addgene 50459-AAV8) or AAV8-hSyn-DIO-hM3D(Gq)-mCherry (Addgene 44361-AAV8). The following stereotaxic coordinates were used to target the CeA in C57BL/6J mice: 1.25 mm posterior from bregma, 2.95 mm lateral to midline, 4.5 mm ventral to skull surface. PbN injections were performed using the following stereotaxic coordinates: 4.9 mm posterior from bregma, 1.2 mm lateral to midline, 3.78 mm ventral to skull surface. To target the CeA in Swiss Webster mice, the following coordinates were used: 1.4 mm posterior from bregma, 3.2 mm lateral to midline and 4.8 mm ventral to skull surface. For PbN injections the following coordinates were used: 5.0 mm posterior to bregma, 1.3 lateral to midline and 3.52 mm ventral to skull surface. Animals recovered for 3 weeks before additional experimental procedures. At the end of the experiments, mice were transcardially perfused with 4% paraformaldehyde solution in 0.1 M Phosphate Buffer (PFA/PB), pH 7.4. Injection sites were verified through histology and only animals with correct injection sites were included in the analyses. For slice electrophysiology experiments, 0.1 µl of rAAV1-hSyn-hChR2(H134R)-EYFP-(Addgene 26973) was injected into the right PbN of *Sst*-cre::Ai9 or *Prkcd*-cre::Ai9 mice. Injections were performed using the following stereotaxic coordinates for *Prkcd*-cre::Ai9: 5.2 mm posterior from bregma, 1.3 mm lateral to midline, 3.52 mm ventral to skull surface. For *Sst*-cre::Ai9 mice coordinates were: 5.1 mm posterior from bregma, 1.2 mm lateral to midline, 3.78 mm ventral to skull surface. Slice electrophysiology experiments were performed at least 4 weeks after viral injection to allow for expression of ChR2 in PbN terminals in the CeA.

### Sciatic cuff implantation

Sciatic nerve surgeries were performed 3 weeks after stereotaxic surgeries as previously described (40, 73). Male and female C57BL/6J or male Swiss Webster mice were randomly assigned to either cuff or sham surgeries such that pairs of co-housed mice underwent the same manipulation. Mice were anesthetized with 2% isoflurane at a flow rate of 0.5 L/min and an incision 1 cm long was made in the proximal one third of the lateral thigh. The sciatic nerve was exposed and gently stretched with forceps inserted under the nerve. The cuff group was implanted with a 2 mm-long-piece of PE-20 non-toxic sterile polyethylene tubing (0.38 mm ID / 1.09 mm OD; Daigger Sci) that was split along its side and slid onto the exposed sciatic nerve. After cuff implantation, the nerve was returned to the thigh. For sham animals, the sciatic nerve was exposed and gently stretched using forceps and then returned to its normal position. The skin was closed with wound clips (Reflex Clips, World Precision Instruments). Mice recovered for at least a one-week before undergoing behavioral testing. All experiments were replicated at least twice.

### Nociceptive testing

Male and female mice were used for nociceptive testing. Testing on males and females were performed separately. Data from both sexes was pooled as no overt sex differences were observed. Individual data points for each sex are clearly identified in all scatter plots. Experimenter was blind to treatment for all behavioral testing and every cohort was counterbalanced to include mice from all experimental groups. Testing was performed on two consecutive days per test. On each testing day, baseline (pre-injection) measurements were taken. Saline or Clozapine-N-oxide (CNO, Enzo Life Sciences, Farmingdale, NY) was injected i.p. (10 mg/kg body weight for hM4Di mice or 3 mg/kg body weight for hM3Dq) and a second measurement (post-injection) was taken 30 minutes to 45 minutes after the i.p. injection. Mice were randomly assigned into control (saline) or experimental (CNO) group on the first day of each test. The next day, the tests were performed with the opposite treatment. The mechanical sensitivity and acetone evaporative tests were performed on the same day, waiting 30 minutes between tests, and 7-8 days after the cuff implantation. Heat sensitivity test was performed 9-10 days after cuff implantation and the Randall-Selitto test for pressure sensitivity on days 11-12. Testing boxes for all tests were 11 × 11 × 13 cm, ventilated and made of opaque Plexiglas.

### Von Frey test

Mice were habituated (for 3 h) to testing chambers placed on an elevated mesh platform prior to behavioral assessment. Von Frey filaments (North Coast Medical, Inc. San Jose,

CA) were used to measure mechanical sensitivity as previously described (22). Starting with the smallest filament, each Von Frey filament, was applied to the mouse hind paws until bent at 30° for ∼2 s. The smallest filament that elicited a paw withdrawal response in at least three of five trials was taken as the mechanical threshold for that trial. The average of 3-5 measurements was calculated individually for each paw and used as the withdrawal threshold.

#### Acetone evaporative test

An adapted acetone evaporative test (74) was used to measure sensitivity to a cold stimulus. An acetone drop was formed at the top of a 1 ml or 3 ml syringe then lightly applied through the wire mesh to the plantar surface of the hind paw ipsilateral (treated) or contralateral (untreated) to sciatic nerve surgery. Following acetone application, nociceptive responses were scored based on responses observed for 60 seconds post application. A modified version of the scoring system described previously for this test (75) was used, with 0 = a rapid transient lifting, licking, or shaking of the hind paw, which subsides immediately; 1 = lifting, licking, and/or shaking of the hind paw, that continues beyond the initial application, but fades within 5 seconds; 2 = protracted, repeated lifting, licking, and/or shaking of the hind paw. The average score of 3-5 stimulations were taken from and used for each hind paw.

#### Hargreaves test

Heat sensitivity in cuff and sham animals was performed using a modified version of the Hargreaves test (76) as previously described (77). Animals were habituated for 1 hour to a Plexiglas testing chamber on an elevated platform with a clear glass surface heated to 30°C. The thermal stimulus was a constant radiant heat source with an active intensity of 25 for C57BL/6J or 32 for Swiss Webster mice directed to the hind paw plantar surface (IITC Life Sciences, Woodland Hills, CA). Active intensity is the intensity of light source as defined by the manufacturer. The time each mouse needed to withdraw the hind paw was recorded. A 15 second cutoff was used to prevent injury. The average of five withdrawal latencies were taken from and used for each hind paw.

#### Randall-Selitto test

A modified Randall-Selitto test (78) was used to measure response thresholds to mechanical pressure stimuli. Male or female mice were lightly anesthetized with 3% isoflurane in an induction chamber, then maintained with 0.5%–1% isoflurane at a flow rate of 0.5 L/min. A sharp pinch not exceeding 200 g of force was delivered to the plantar surface of the paw ipsilateral and contralateral to cuff or sham implanted sciatic nerve. Pinch pressure for withdrawal was recorded at 1-minute intervals for 30 minutes. The average of five trials was calculated individually for each animal.

#### Formalin test

Male and female C57BL/6J or male Swiss Webster mice were habituated for 1 hour in plexiglass testing chambers on an elevated platform with transparent floors. A mirror was positioned directly below the chambers to properly visualize mice hind paws. After habituation, mice received i.p. injection of either CNO (10mg/kg) or saline and were immediately returned to the testing chamber. The experimenter was blind to treatment. 30-40 minutes after i.p. injection, C57BL/6J mice were injected with 10 μl of 2-3% formalin and Swiss Webster with 10 μl of 5% formalin into either left or right hind paws. Immediately after, they were returned to the testing chambers and time spent in nociceptive behaviors, defined as licking, lifting and shaking the hind paws, were individually measured for 40 minutes (C57BL/6J) or 60 minutes (Swiss Webster) in 5 minutes intervals. Total time spent in spontaneous nociceptive behaviors was defined as the sum of the time spent in the individual behaviors. Phase 1 of the formalin test was defined as the first five minutes post-formalin injection and phase 2 was measured from 5 to 40 minutes after formalin injection.

### Immunohistochemistry

At the end of the experiments, mice were deeply anesthetized with 1.25% Avertin anesthesia (2,2,2-tribromoethanol and tert-amyl alcohol in 0.9% NaCl; 0.025 ml/g body weight), then perfused transcardially with 0.9% NaCl (37°C), followed by 100 mL of ice-cold 4% paraformaldehyde in phosphate buffer (PFA/PB). Immediately after perfusion, we dissected the brains, post fixed in 4% PFA/PB overnight at 4°C and cryoprotected in 30% sucrose/PB for 48 hours. Thirty μm coronal sections containing the regions of interest (central amygdala and/or parabrachial nucleus) were collected in 0.1 M Phosphate Buffered Saline (PBS), pH 7.4 containing 0.01% sodium azide (Sigma) using a freezing sliding microtome. Sections were stored in 0.1 M Phosphate Buffered Saline (PBS), pH 7.4 containing 0.01% sodium azide (Sigma) at 4°C until used for immunostaining. After rinsing in PBS, sections were incubated in 0.1% Triton X-100 in PBS for 10 minutes at room temperature and were then blocked in 5% normal goat serum (NGS) (Vector Labs, Burlingame, CA) with 0.1% Triton X-100, 0.05% Tween-20 and 1% bovine serum albumin (BSA) for 30 minutes at room temperature. Primary antibody incubations were for overnight or 72 hours at 4°C, followed by 1 hour at room temperature. Sections were then rinsed in PBS and incubated in secondary antibodies in 1.5% NGS blocking solution with 0.1% Triton X-100, 0.05% Tween 20 and 1% BSA, protected from light, for 2 hours at room temperature. Sections were then rinsed in PBS, mounted on positively charged glass slides and air-dried overnight. Coverslips were placed using DAPI Fluoromount-G mounting media (Southern Biotech) and slides were stored at room temperature overnight and then stored under 4°C.

#### Verification of brain injection sites for behavioral experiments

For injection site verification, the following primary and secondary antibodies were used: rat anti-mCherry (1:250 for 72 hours or 1:125 overnight, Invitrogen, M11217), chicken anti-GFP (1:1000 for 72 hours or 1:500 for overnight, Invitrogen, ab13970), rabbit anti-β-gal (1:1000, Millipore Sigma, ab986 for 72 hours or 1:500 overnight) and goat anti-rat Cy3 (1:250, Invitrogen, A10522), Alexa Fluor 647-conjugated goat anti-rabbit (1:500, Invitrogen, A21244) and/or Alexa Fluor 488-conjugated goat anti-chicken IgY (H+L) (1:1000, Invitrogen, A-11039). Correct injection sites were defined as brains with mCherry+ cells localized to the PbN and mCherry+ terminals and LacZ transduced cell bodies localized to the CeA.

#### c-Fos monitoring

For c-Fos experiments to validate CNO-mediated activation of neurons expressing hM3dq, mice were habituated for 1 hour in a behavioral room with red lights. Mice then received i.p. injection of either CNO (3mg/kg) or saline and were immediately returned to their cages. 30-35 minutes after saline or CNO i.p. injections, animals were then anesthetized with 1.25% Avertin i.p. (0.4 mg/g) injection and perfused as previously described. For pinch-induced c-Fos experiments, male and female mice were euthanized and perfused as described above 1 hour after the completion of the Randall-Selitto test. For both c-Fos experiments, the following primary antibodies were used: rat anti-mCherry (1:250 for 72 hours or 1:125 overnight, Invitrogen, M11217) and rabbit anti-Phospho-c-Fos (Ser32) (1:2000 for 72 hours or 1:1000 overnight, Cell Signaling Technology, #5348). For secondary antibodies: goat anti-rat Cy3 (1:250, Invitrogen, A10522) and Alexa Fluor 647-conjugated goat anti-rabbit (1:500, Invitrogen, A21244) were used.

### Immunohistochemistry to verify injection sites for electrophysiological experiments

Slices were fixed in 4% PFA at 4 °C for 48 hours at the end of recordings. Following slice fixation, slices containing the CeA and PbN were stored in 0.1% sodium azide in PBS 4 °C until histological processing. Slices were rinsed in 0.1 M PBS and incubated in 0.1% Triton-X-100 in PBS for 10 minutes at room temperature. 5% NGS-based blocking buffer (0.1% Triton-X-100, 0.05% Tween-20, and 1% BSA) was used for slice incubation at room temperature for 30 minutes. Slices were then rinsed in PBS and incubated in 1.5% NGS-based blocking buffer (0.1% Trition-X-100, 0.05% Tween-20, and 1% BSA) containing the primary antibody rabbit anti-GFP (Invitrogen, A6455, 1:250 concentration) for 1 week at 4 °C followed by 1 hour at room temperature. Slices were then rinsed in PBS and incubated in 1.5% NGS-based blocking buffer (0.1% Trition-X-100, 0.05% Tween-20, 1% BSA) containing the secondary antibody goat anti-Rabbit IgG (H+L) Highly Cross-Adsorbed (Alexa Fluor 488, Invitrogen A11034, 1:100 concentration) overnight at room temperature under minimal light. Following secondary antibody incubation, slices were rinsed in PBS and incubated for 10 minutes in a series of increasing concentrations of 2,2’-thiodiethanol (TDE, Sigma) to allow for tissue clearing prior to image acquisition (79) in the following order of concentrations of TDE in PBS: 10%, 30%, 60%, and 80% followed by a 2-hour incubation in 97% TDE solution. All slice incubations in TDE were at room temperature. Slices were mounted on positively charged glass slides, coverslips were placed on slices covered with 97% TDE, and slides were sealed with either clear nail polish or Fluoromount-G mounting media (SouthernBiotech).

For optogenetic assisted circuit mapping, correct injection sites were defined as brains with localized GFP-expressing cell bodies at the PbN and GFP+ terminals at the CeA. For electrophysiological experiments to validate intersectional approach, correct injection sites were defined as brains with GFP-expressing transduced cells at the CeA, mCherry+ transduced cells at the PbN and mCherry-expressing terminals at the CeA.

### Image acquisition and analysis

Images were acquired using a Nikon A1R laser scanning confocal microscope or a Leica DM5500 using a 2X (for low magnification) or a 10X (for high magnification) objective. 40X oil-immersion objective was used for high magnification representative images. Experimenter was blind to experimental group and analyses were performed on images collected using a 10X objective. GFP, RFP and CY5 channels were used for consecutive image acquisition, and z stack images were collected at 5.4 μm steps. Imaging parameters (laser intensity, gain, and pinhole) were kept identical between experiments. NIS Elements software automatically stitched acquired images and converted the stacks into maximum intensity z-projections. Once images were acquired, rostro-caudal level and anatomical location of positive cells were determined using distinctive anatomical landmarks and a mouse brain atlas (80). Mapping of injection sites were delineated based on the anatomical regions containing transduced cells.

#### Quantification of positive cells

Quantitative analyses were performed between rostro-caudal levels −0.94 and −1.70 relative to bregma for CeA and between −4.96 and −5.34 relative to bregma for PbN. Anatomical delineation for each brain region was determined using a mouse brain atlas (Paxinos and Franklin, 2001). Quantification of positive cells was performed using the NIS Elements software in each channel on one section per rostro-caudal level per mouse. Labeled and co-labeled cells were automatically identified with the NIS Elements software and were visually confirmed by an experimenter. Total numbers of c-Fos and mCherry + cells were defined as the sum of positive cells in 4 PbN slices between −4.96 and −5.34 relative to bregma per animal.

### *Ex-vivo* electrophysiology

#### Acute CeA and PbN slice preparations

To confirm the effects of clozapine N-oxide (CNO) on hM4Di-transduced cells in the PbN, acute PbN slices were prepared from male C57BL/6J mice from 12 to 18 weeks stereotaxically injected with AAV.hSyn.HI.eGFP-Cre into the CeA and AAV8-hSyn-DIO-hMD4i-mCherry into the PbN. To validate a functional circuit between the PbN and CeA, acute CeA slices were prepared from either *Prkcd*-Cre::Ai9 or *Sst*-Cre::Ai9 male mice (25-26 weeks) injected with rAAV1-hSyn-hChR2(H134R)-EYFP. All electrophysiological experiments were performed at least 4 weeks after stereotaxic injections to allow for efficient uptake of retrograde virus in the PbN (for chemogenetic validation experiments) or efficient expression of ChR2 in PbN terminals in the CeA (for opto-assisted circuit mapping).

Mice were deeply anesthetized with 1.25% Avertin i.p. (0.4 mg/g) and transcardially perfused with an ice-cold cutting solution including: 110 mM choline chloride, 25 mM NaHCO_3_, 1.25 mM NaH_2_PO_4_, 2.5 mM KCl, 0.5 mM CaCl_2_, 25 mM D-glucose, 12.7 mM L-ascorbic acid, and 3.1 mM pyruvic acid. Brains were immediately dissected and submerged in ice-cold cutting solution. Coronal sections including the right CeA (250 µm) and PbN (300 µm) were cut using feather carbon steel blades (Ted Pella Inc., Redding, CA) and a Leica VT1200 S vibrating blade microtome (Leica Microsystems Inc., Buffalo Grove, IL). CeA and PbN slices were transferred into a recovery chamber of oxygenated artificial cerebrospinal fluid (ACSF) containing 125 mM NaCl, 2.5 mM KCl, 1.25 mM NaH_2_PO_4_, 25 mM NaHCO_3_, 2.0 CaCl_2_, 1.0 MgCl_2_, and 25 mM D-glucose. Slices recovered for 30 minutes at 33 °C and were then transferred to room temperature for at least 20 minutes prior to recordings. Cutting solution and ACSF were continuously saturated with 95%/5% O_2_/CO_2_.

#### Electrophysiological Recordings

Whole-cell patch-clamp recordings of neurons located in either the PbN or CeA were collected at 33 ± 1 °C. The recording chamber was perfused with oxygenated ACSF (95%/5% O_2_/CO_2_) at a flow rate of 1 mL/min. An in-line solution heater and heated recording chamber (Warner Instruments) monitored and controlled the temperature conditions throughout the recordings. Neurons were identified through differential interference contrast optics and fluorescent microscopy using an upright telescope (Nikon Eclipse FN1). Recording pipettes (2.5-5 MΩ resistance for CeA and 4-7 MΩ resistance for PbN) were filled with a potassium methyl sulfate-based internal solution (120 mM KMeSO_4_, 20 mM KCl, 10 mM HEPES, 0.2 mM EGTA, 8 mM NaCl, 4 mM Mg-ATP, 0.3 Tris-GTP, and 14 mM phosphocreatine, with pH adjusted to 7.3 with KOH (approximately 300 mosmol-1)). Recordings were performed using a Multiclamp 700B patch-clamp amplifier interfaced with a Digidata 1550 acquisition system and pCLAMP 10.7 software (Molecular Devices) installed on a Dell computer.

#### Channelrhodopsin-2 (ChR2)-assisted circuit mapping

Optically evoked excitatory postsynaptic currents (oEPSCs) of tdTomato-expressing PKCδ+ or Som+ CeA neurons were recorded at a holding potential of -70 mV, and a single light pulse of 10 ms duration, delivered at 10 Hz, was presented to elicit oEPSCs. Signals were filtered at 10 kHz and acquired at 100 kHz. A blue LED illumination system (λ = 470 nm, Mightex) was used to stimulate PbN-projecting terminals within the CeA. An optical power console and sensor (Thorlabs) measured the blue light output directly through the 40x objective of the microscope prior to each experiment, and the intensity output measured between 10-12 mW. Prior to formation of membrane to pipette seals (∼1 GΩ), tip potentials were zeroed. Additionally, the pipette capacitances were compensated and series resistances (not exceeding 20 MΩ) were repeatedly monitored throughout the duration of the recording. Whole-cell capacitance recordings under the voltage-clamp configuration were collected at a holding potential of -70mV and then presented a ± 10 mV voltage change of 25 ms duration. Under current-clamp configuration, 500 ms depolarizing current injections of 220 and 280 pA amplitudes were used to elicit repetitive action potential firing. Cells that categorized as late-firing or regular-spiking based on their latencies to fire in response to prolonged depolarizing current injections, as previously defined (33). Briefly, regular-spiking (69) neurons are categorized if the latency to the first spike is shorter than 100 ms and late-firing (37) neurons are categorized if the latency to the first spike is longer than 100 ms. Spike latency for RS and LF cells were assessed in response to 220-pA current step for Som+ neurons and in response to 280-pA current step for PKCδ+ neurons, which elicited an average of 10 spikes in recorded neurons. Current clamp signaling was filtered at 10 kHz and acquired at 100 kHz. The anatomical location of each CeA neuron recorded was determined using a mouse brain atlas (80).

#### Electrophysiological validation of chemogenetic approach

Current-clamp recordings were performed on hM4Di-transduced PbN cells to assess neuronal excitability before and after CNO bath application. All recordings were performed in ACSF with the addition of synaptic blockers (5 μM CPP (-((R)-2-carboxypiperazin-4-yl)-propyl-1-phosphonic acid), 10 μM NBQX (2,3-dioxo-6-nitro-1,2,3,4-tetrahydrobenzo[f]quinoxaline-7-sulfonamide) and 5 μM GABAzine (6-imino-3-(4-methoxyphenyl)-1(6H) pyridazinebutanoic acid hydrobromide). PbN cells expressing mCherry were visually identified and targeted, and series resistances of assessed neurons did not exceed 40 MΩ. To account for the heterogenous firing phenotypes observed within the PbN (i.e. spontaneously active, low threshold, regular-spiking, or late-firing neurons), current clamp protocols were conducted respective to the firing properties. Spontaneously active neurons were recorded continuously for 20 minutes, while low threshold neurons were recorded using an 800 ms depolarizing current ramp protocol. Action potential firing in regular-spiking or late-firing PbN neurons were recorded in response to a 500 ms square pulse depolarizing current injections. Current injection amplitudes that elicit between 2-5 stable action potentials were used every 15 seconds. At least 5 stable recordings were obtained prior to bath application of either 10 µM CNO or vehicle (i.e. saline) in ACSF. The number of elicited action potentials during each condition were averaged across 5 traces to assess the effect of CNO on excitability. Current clamp signals were acquired at 100 kHz and filtered at 10 kHz.

### Statistical analysis

Data are presented as mean ± SEM. Statistical analysis was performed using unpaired or paired two tailed t test, or two-way analysis of variance (ANOVA) followed by Tukey’s multiple comparison tests using Graph Pad Prism version 9.0. The significance level was set at p < 0.05. Sample sizes and p values are described in each figure legend. Detailed information on each statistical test performed are shown in **Supplemental Table 1**.

## Acknowledgments

This research was supported by the National Center for Complementary and Integrative Health Intramural Research Program (YC), a fellowship from the NIH Postdoctoral Research Associate Training (OSC), and a fellowship from the NIH Center on Compulsive Behaviors (SS). We would like to thank Dr. Maria Luisa Torruella-Suarez for feedback on the manuscript and project, Barbara Benowitz, Benjamin Neugebauer and Adela Francis Malavé for technical assistance in the project, and the National Institutes of Neurological Disorder Mouse Facility staff for their vital work in animal husbandry.

## Notes

### Competing Interest Statement

The authors have declared no competing interest.

